# A minimal CRISPR-Cas3 system for genome engineering

**DOI:** 10.1101/860999

**Authors:** Bálint Csörgő, Lina M. León, Ilea J. Chau-Ly, Alejandro Vasquez-Rifo, Joel D. Berry, Caroline Mahendra, Emily D. Crawford, Jennifer D. Lewis, Joseph Bondy-Denomy

## Abstract

CRISPR-Cas technologies have provided programmable gene editing tools that have revolutionized research. The leading CRISPR-Cas9 and Cas12a enzymes are ideal for programmed genetic manipulation, however, they are limited for genome-scale interventions. Here, we utilized a Cas3-based system featuring a processive nuclease, expressed endogenously or heterologously, for genome engineering purposes. Using an optimized and minimal CRISPR-Cas3 system (Type I-C) programmed with a single crRNA, large deletions ranging from 7 - 424 kb were generated in *Pseudomonas aeruginosa* with high efficiency and speed. By comparison, Cas9 yielded small deletions and point mutations. Cas3-generated deletion boundaries were variable in the absence of a homology-directed repair (HDR) template, and successfully and efficiently specified when present. The minimal Cas3 system is also portable; large deletions were induced with high efficiency in *Pseudomonas syringae* and *Escherichia coli* using an “all-in-one” vector. Notably, Cas3 generated bi-directional deletions originating from the programmed cut site, which was exploited to iteratively reduce a *P. aeruginosa* genome by 837 kb (13.5%) using 10 distinct crRNAs. We also demonstrate the utility of endogenous Cas3 systems (Type I-C and I-F) and develop an “anti-anti-CRISPR” strategy to circumvent endogenous CRISPR-Cas inhibitor proteins. CRISPR-Cas3 could facilitate rapid strain manipulation for synthetic biological and metabolic engineering purposes, genome minimization, and the analysis of large regions of unknown function.

## Introduction

CRISPR-Cas systems are a diverse group of RNA-guided nucleases^1^ that defend prokaryotes against viral invaders^2,3^. Due to their relatively simple architecture, gene-editing applications have focused on Class 2 CRISPR systems^4^ (i.e. Cas9 and Cas12a), but Class 1 systems hold great potential for gene editing technologies, despite being more complex^5–7^. The signature gene in Class 1 Type I systems is Cas3, a 3’-5’ ssDNA helicase-nuclease enzyme that, unlike Cas9 or Cas12a, degrades target DNA processively^5,6,8–13^. This property of CRISPR-Cas3 systems raises the possibility of its development as a tool for large genomic deletions.

Organisms from all domains of life contain vast segments of DNA that are poorly characterized, of unknown function, or may be detrimental for fitness. In prokaryotes, these include prophages, plasmids, and mobile islands, while in eukaryotes, large regions of repetitive sequences and non-coding DNA are poorly characterized. Methods for generating programmable and rapid large genomic deletions are needed for the study and engineering of these regions, however these are currently inefficient^14^. A methodology that allows for targeted large genomic deletions in any host, either with precisely programmed or random boundaries, would be broadly useful^15^.

Type I systems are the most prevalent CRISPR-Cas systems in nature^1^, which has enabled the use of endogenous CRISPR-Cas3 systems for genetic manipulation via self-targeting. This has been accomplished in *Pectobacterium atrosepticum* (Type I-F)^16^, *Sulfolobus islandicus* (Type I-A)^17^, in various *Clostridium* species (Type I-B)^18–20^, and in *Lactobacillus crispatus* (Type I-E)^21^, leading to deletions as large as 97 kb amongst the self-targeted survivor cells^16^. Additionally, two recent studies have repurposed Type I systems for use in human cells, including the ribonucleoprotein (RNP) based delivery of a Type I-E system^22^, and the use of I-E and I-B systems for transcriptional modulation^23^. Here, we repurposed a Type I-C CRISPR system from *Pseudomonas aeruginosa* for genome engineering in microbes. Importantly, by targeting the genome with a single crRNA and selecting only for survival after editing, this tool is a counter-selection-free approach to programmable genome editing. CRISPR-Cas3 is capable of efficient genome-scale modifications currently not achievable using other methodologies. It has the potential to serve as a powerful tool for basic research, discovery, and strain optimization.

## Results

### Implementation and optimization of genome editing with CRISPR-Cas3

Type I-C CRISPR-Cas systems utilize just three *cas* genes to produce the crRNA-guided Cascade surveillance complex that can recruit Cas3: *cas5, cas8*, and *cas7* (Figure 1A), making it a minimal system^24,25^. A previously constructed *Pseudomonas aeruginosa* PAO1 strain (PAO1^IC^) with inducible *cas* genes and crRNAs^26^ was used here to conduct targeted genome manipulation. The expression of a crRNA targeting the genome caused a transient growth delay (Figure 1B), but survivors were isolated after extended growth. By targeting *phzM*, a gene required for production of a blue-green pigment (pyocyanin), we observed yellow cultures (Figure 1C) for 16 out of 36 (44%) biological replicates (18 recovered isolates from two independent *phzM* targeting crRNAs). PCR of genomic DNA confirmed that the yellow cultures had lost this region, while blue-green survivors maintained it (Supplementary Figure 1). Three of these PAO1^IC^ deletion strains were sequenced, revealing deletions of 23.5 kb, 52.8 kb, and 60.1 kb, and each one was bi-directional relative to the crRNA target site (Figure 1D). This demonstrated the potential for Type I-C Cas3 systems to be used to induce large genomic deletions with random boundaries surrounding a programmed target site.

**Figure 1.**
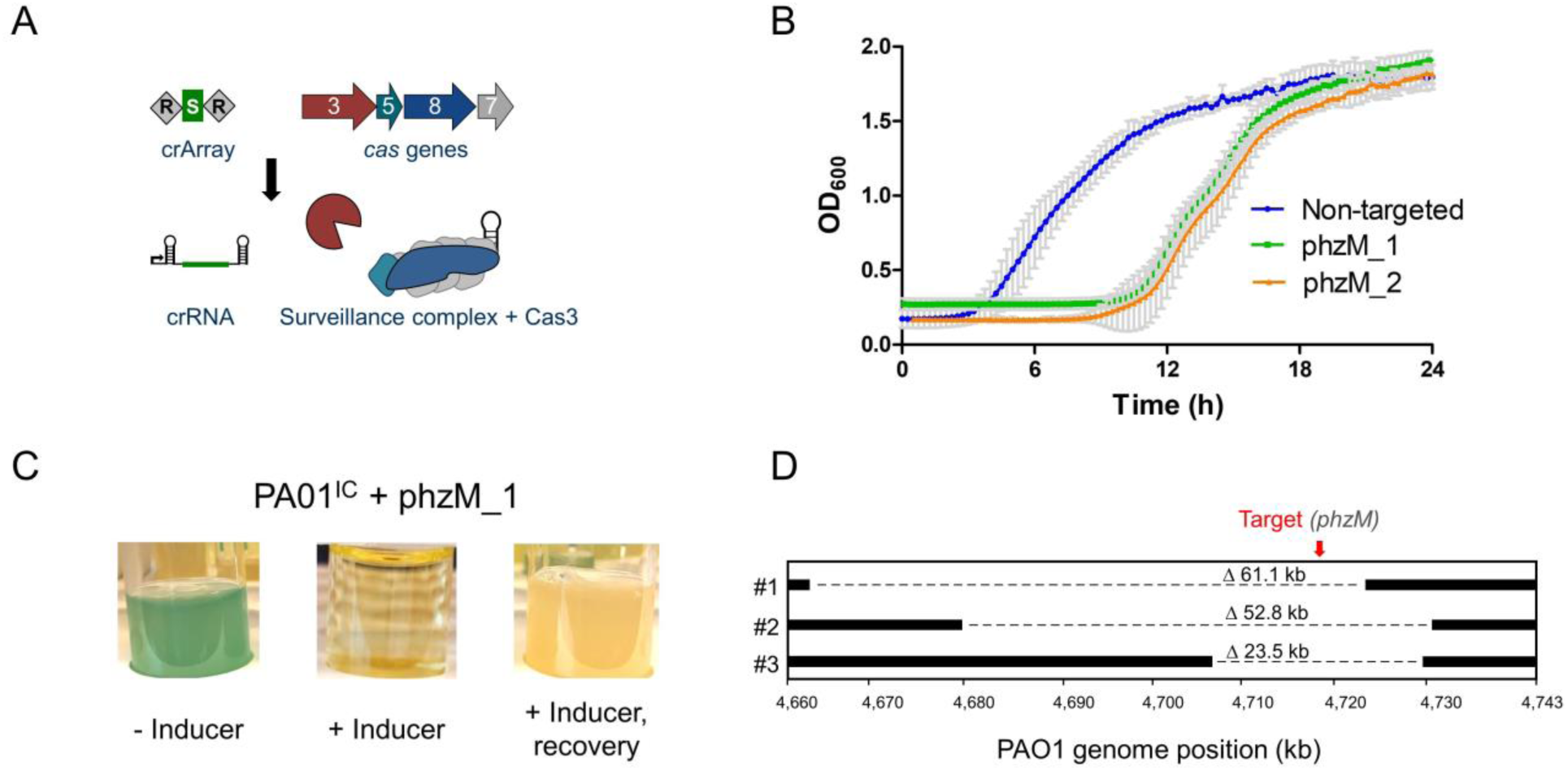
**A)** A schematic of the Type I-C *cas* gene operon and CRISPR array. The surveillance complex is made up of Cas proteins (Cas5_1_:Cas8_1_:Cas7_7_) and one crRNA, which recruits Cas3 upon target DNA recognition. **B)** Growth curves of 2 PAO1^IC^ strains expressing different crRNAs targeting *phzM* (green and orange) compared to a non-targeting strain (blue). Values are the mean of 8 biological replicates each, error bars indicate SD values. **C)** Cultures resulting from *phzM* targeting, in the absence of inducer (-ind), presence (+ind), and after recovery. **D)** Whole-genome sequencing of three PAO1^IC^ self-targeted survivor strains. Bars indicate boundaries of deletions; red arrow indicates genomic position of targeted sequences.

To determine the *in vivo* processivity of the Cas3 enzyme, we targeted 2 of the 16 e<u>x</u>tended <u>n</u>on-<u>es</u>sential (XNES) regions >100 kb in length (Supplementary Table 1) identified from a transposon sequencing (TnSeq) data set^27^. The frequency of deletions generated by crRNAs targeting XNES 1 and XNES 2 (along with additional targeting of *phzM*, which is found in XNES 15) was quantified revealing that 20-40 % of the surviving colonies had deletions (Figure 2A). To understand how cells lacking large deletions had survived self-targeting, three possibilities were considered: i) a *cas* gene mutation, ii) a PAM or protospacer mutation, or iii) a mutation to the plasmid expressing the crRNA. Three non-deletion survivors from each of the six self-targeting crRNAs had functional *cas* genes when the self-targeting crRNA was replaced with a phage targeting crRNA (Supplementary Figure 2A), and target sequencing revealed no mutations. PCR-amplification and sequencing of the crRNA-expressing plasmids isolated from the survivors revealed the primary escape mechanism: recombination between the direct repeats, leading to the loss of the spacer (Supplementary Figure 2B). The resulting bands from an additional 17 survivors that lacked deletions were also ∼60 bp shorter (Supplementary Figure 2C), consistent with the loss of one repeat and spacer.

**Figure 2.**
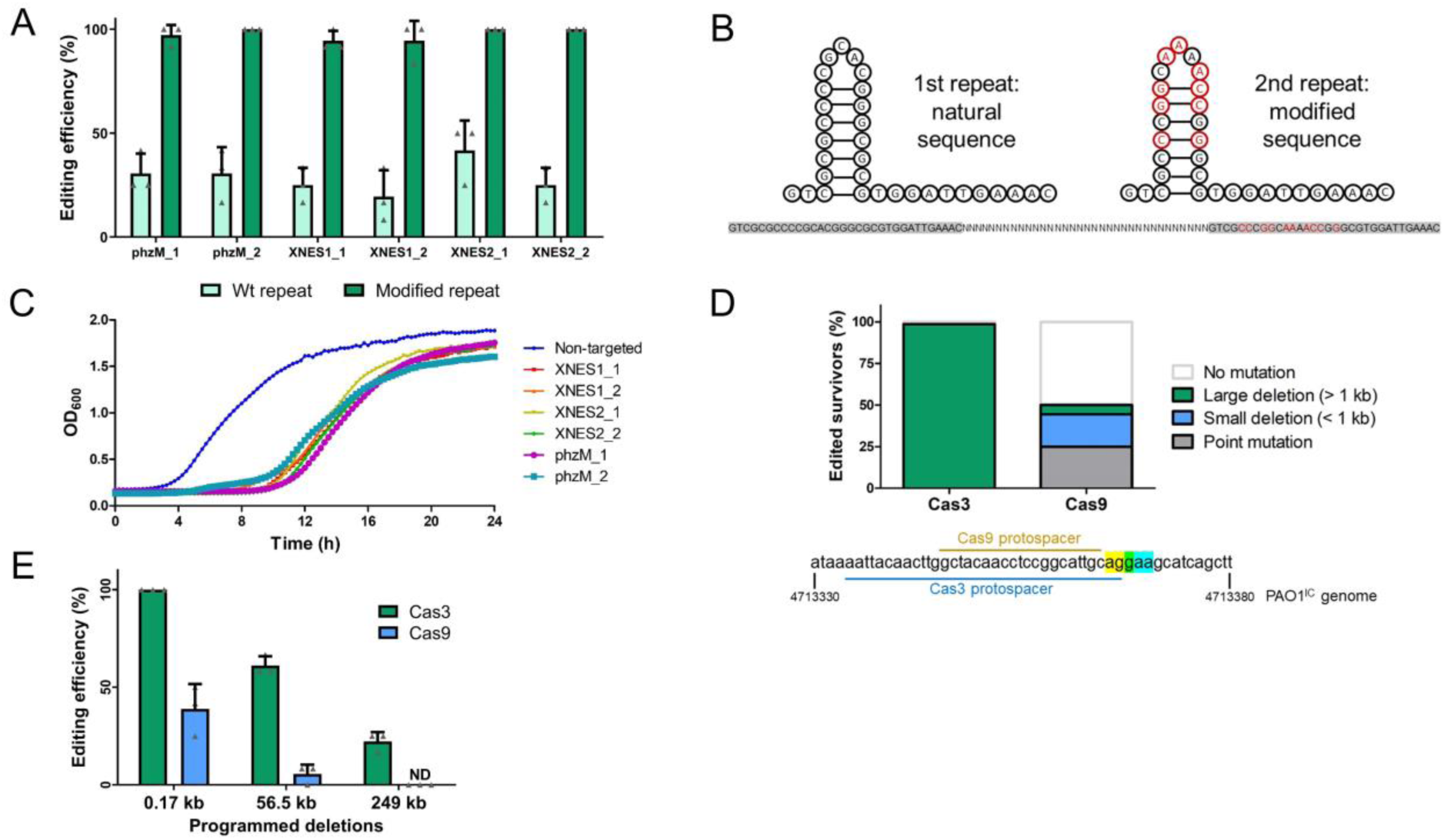
**A)** Percentage of survivors with a genomic deletion at the location targeted. Six different crRNA constructs with either wild-type (Wt) repeat sequences (light green) or with the second repeat being modified (dark green). Values are means of 3 biological replicates each, where 12 individual surviving colonies were assayed per replicate, error bars show SD values. **B)** Sequence and structure of natural and modified repeat sequences. Specifically engineered modified nucleotides shown in red; repeat sequences highlighted in gray with an arbitrary intervening spacer sequence. **C)** Growth curves of PAO1^IC^ strains expressing distinct self-targeting crRNAs flanked by modified repeats. Non-targeting crRNA expressing control is marked in blue. Values depicted are averages of 4 biological replicates each. **D)** Gene editing outcomes for distinct survivor cells targeted with either a Type II-A SpyCas9 system or a Type I-C Cas3 system (n=72). **E)** Percentage of survivors with the specific deletion size present (0.17 kb, 56.5 kb, or 249 kb) using homologous repair templates with the Cas3 system (green) or the SpyCas9 system (blue). Values are means of 3 biological replicates each, where 12 individual surviving colonies were assayed per replicate, error bars show SD values, ND: not detected.

Spacer excision was successfully prevented by engineering a modified repeat (MR), with six mutated nucleotides in the stem and three in the loop of the second repeat (Figure 2B), disrupting homology between the two direct repeats. A phage-targeting crRNA with this new design targeted phage as well or better than the same crRNA with unmodified repeats (Supplementary Figure 3A). Using the same self-targeting spacers designed against *phzM*, XNES 1, and XNES 2 with the MR resulted in a robust increase in editing efficiencies to 94-100% for the six tested crRNAs (Figure 2A) and spacer excision was no longer detected. 211 of 216 (98 %) total survivor cells had large deletions based on PCR screening (i.e. > 1 kb), while the remaining 5 had inactive CRISPR-Cas systems when tested with the phage-targeting crRNA (Supplementary Figure 3B).

The processivity of Cas3 could likely lead to unintended deletions of neighboring essential genes, if targeting is initiated nearby. To assess the phenotype of such an event, we intentionally targeted an essential gene, *rplQ* (a 50S ribosomal subunit protein^28^). Two different MR crRNAs targeting *rplQ* led to a severely extended lag time compared to non-essential gene targeting. Only 8 out of 36 *rplQ*-targeting biological replicates grew after 24 hours, compared to the transient growth delay of ∼12 hours when targeting non-essential genes (Supplementary Figure 4A). Subsequent analysis of these 8 survivor cultures with phage targeting assays revealed non-functional *cas* genes (Supplementary Figure 4B). Importantly, no spacer excision events were detected in this experiment or among the 216 replicates screened above. This experiment highlights the robustness of the deletion method, as the outcome of essential gene versus non-essential gene targeting is noticeably distinct.

### Comparison of Cas3 and Cas9 based editing

To determine whether large deletions are a direct consequence of the Cas3 enzyme, we compared self-targeting outcomes to an isogenic strain expressing the non-processive *Streptococcus pyogenes* Cas9, PAO1^IIA^. Two crRNAs (as sgRNAs) that partially overlapped with the crRNAs used for PAO1^IC^ were targeted to *phzM* (Figure 2E, Supplementary Figure 5). PCR and sequencing analysis of these surviving cells revealed that deletions larger than 1 kb are a rare occurrence (5.6 % assayed survivor cells, n = 72) compared to 98.6 % with PAO1^IC^ (Figure 2E). The more common modes of survival were small deletions between 0.2 – 0.5 kb in length (19.4 % of all survivors), or 1-3 bp protospacer/PAM deletions/mutations (25 %). Whole-genome sequencing (WGS) of two large deletion survivors selected for by Cas9 showed lesions of 5 kb and 23 kb around the target site, respectively. The frequent isolation of small deletions from targeting with the non-processive SpyCas9 directly implicates Cas3’s enzymatic activity as the cause of large deletions.

The direct relationship between Cas3 nuclease-helicase activity and survival via large deletions led us to hypothesize that its processive ssDNA nuclease activity may promote recombination. To test this, we provided a repair template with ∼500 bp of the upstream and downstream regions flanking the desired deletion to enable homology directed repair (HDR). We chose 0.17 kb and 56.5 kb deletions around *phzM* and a 249 kb deletion within XNES8 for the programmed deletions (Supplementary Figure 6). The recombination efficiencies were significantly higher with Cas3 than with Cas9 (Figure 2F). The 249 kb deletion was incorporated in 22 % of the Cas3-generated survivors, compared to 0% using Cas9 (*χ*^*2*^ (1, N = 72) = 9, *p* = 2.7E-03). The 56.5 kb deletion had an efficiency of 61 % vs. 5.5 % (*χ*^*2*^ (1, N = 72) = 25, *p* = 5.73E-07), and the 0.17 kb deletion had an efficiency of 100 % vs. 39 % when targeting with Cas3 or Cas9, respectively (*χ*^*2*^ (1, N = 72) = 31.68, *p* = 1.82E-08). These data support the hypothesis that Cas3 enhances recombination at cleavage sites.

### Rapid genome minimization of P. aeruginosa using CRISPR-Cas3 editing

Large deletions with undefined boundaries provide an unbiased mechanism for genome streamlining, screening, and functional genomics. To demonstrate the potential for Cas3, we aimed to minimize the genome of *P. aeruginosa* through a series of iterative deletions of the XNES regions (Figure 3A). Six XNES regions (including XNES 15, carrying *phzM*) were iteratively targeted in six parallel lineages (Figure 3B), resulting in 35 independent deletions (WGS revealed no deletion at XNES 2 in one of the strains). Deletion efficiency remained high (> 80 %) throughout each round of self-targeting (Supplementary Figure 7). WGS of these 6 multiple deletion strains (*Δ6*_*1*_ -*Δ6*_*6*_) revealed that no two deletions had the exact same coordinates, highlighting the stochastic nature of the process. The smallest isolated deletion was 7 kb and the largest 424 kb (mean: 92.9 kb, median: 58.2 kb). Of note, 4 genes (PA0123, PA1969, PA2024, and PA2156) previously identified as essential^27^ were deleted in at least one of the lineages. Most deletions appeared to be resolved by flanking microhomology regions (Supplementary Table 2), implicating alternative-end joining^29^ as the dominant repair process.

**Figure 3.**
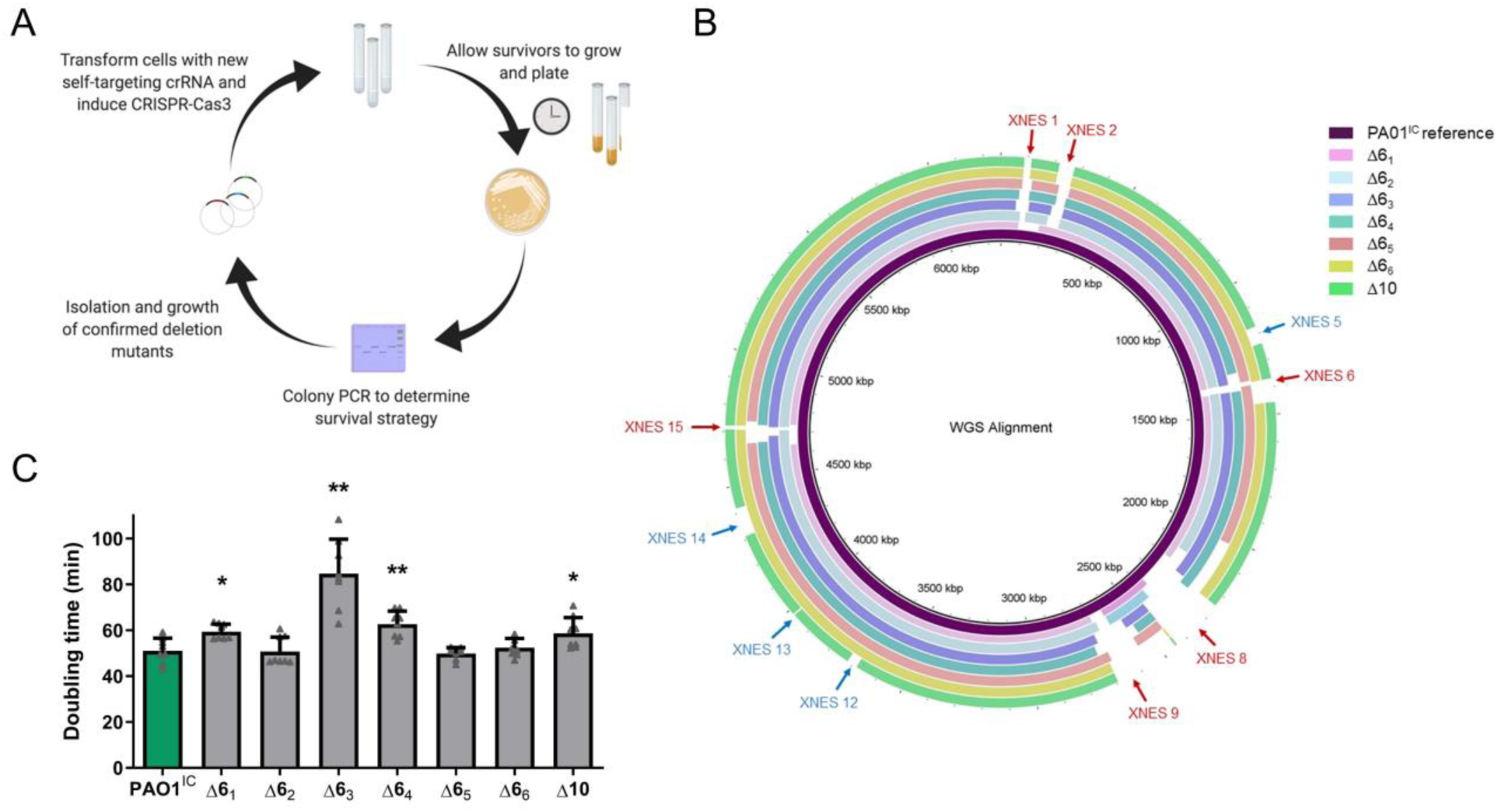
**A)** Schematic overview of the iterative deletion generating process. **B)** Whole-genome sequences of six PAO1^IC^ strains that have been iteratively targeted at six distinct genomic positions and one (derived from strain Δ6_6_) with ten total deletions (Δ10) aligned to the parental *P. aeruginosa* PAO1^IC^ strain. The first six targeted sites are marked with red arrows, and the final four are marked with blue arrows. **C)** Calculated doubling times of the seven genome-reduced strains (strains Δ6_1_ – 6_6_ with six deletions, Δ10 with ten) compared to the parent PAO1^IC^ strain (green). Values are means of 8 biological replicates, error bars represent SD values, * p < 0.05, ** p < 0.01, paired T-test compared to PAO1^IC^.

To minimize the genome further, one of the already reduced strains was subjected to 4 additional rounds of deletions at XNES regions for a total of 10 genomic deletions (*Δ10*, Figure 3B). Whole-genome sequencing of the *Δ10* strain showed a genome reduction of 849 kb (13.55 % of the genome). Generation of large deletions resulted in a growth defect in some cases, with significantly slower growth in 3 of the 6 deletions strains (Δ6_1_, Δ6_3_, and Δ6_4_), with the other 3 growing normally (Figure 3C). *Δ10* also displayed a decrease in fitness, showing a ∼15 % increase in doubling time compared to the parent strain. The general subtlety of the growth defects was likely bolstered by the selection of fast-growing colonies at each round of deletion.

### CRISPR-Cas3 editing in distinct bacteria

To enable expression of this system in other hosts, we constructed an all-in-one vector (pCas3cRh) carrying the I-C specific crRNA with a modified repeat sequence and *cas3, cas5, cas8*, and *cas7* (Supplementary Figure 8A). As a pilot experiment, we transformed wild-type PAO1 with a non-targeting crRNA and crRNAs targeting *phzM* and *XNES2*. Induction of the targeting crRNAs induced editing efficiencies between 95-100 % (Supplementary Figure 8B-D).

Having verified that pCas3cRh was functional, we tested this system in the model organism *Escherichia coli* K-12 MG1655. crRNAs were designed to target *lacZ* or its vicinity (Figure 4A), where it is flanked by non-essential DNA (124.5 kb upstream, 22.4 kb downstream). Transformations were plated directly on inducing media containing X-gal and scored using blue/white screening. Depending on the crRNA used, directly targeting *lacZ* or 30 kb upstream yielded 51-90% or 82-85% editing efficiencies, respectively (Figure 4B). 95 of the 96 LacZ (-) survivors assayed by PCR showed an absence of the *lacZ* region. crRNAs downstream of *lacZ*, however, had reduced efficiency as they approached the essential gene, *hemB. frmA* targeting (13 kb downstream of *lacZ*) had lower editing efficiencies (21-25 %) and *yaiS* (18 kb downstream of *lacZ*) even lower (2%). This drop in efficiency was independent of the strand being targeted (and therefore the predicted strand for Cas3 loading and 3’-5’ translocation), confirming the importance of Cas3 bi-directional deletions. Indeed, WGS of selected Δ*lacZ* cells revealed bi-directional deletions ranging from 17.5-106 kb encompassing the targeted region (Figure 4C).

**Figure 4.**
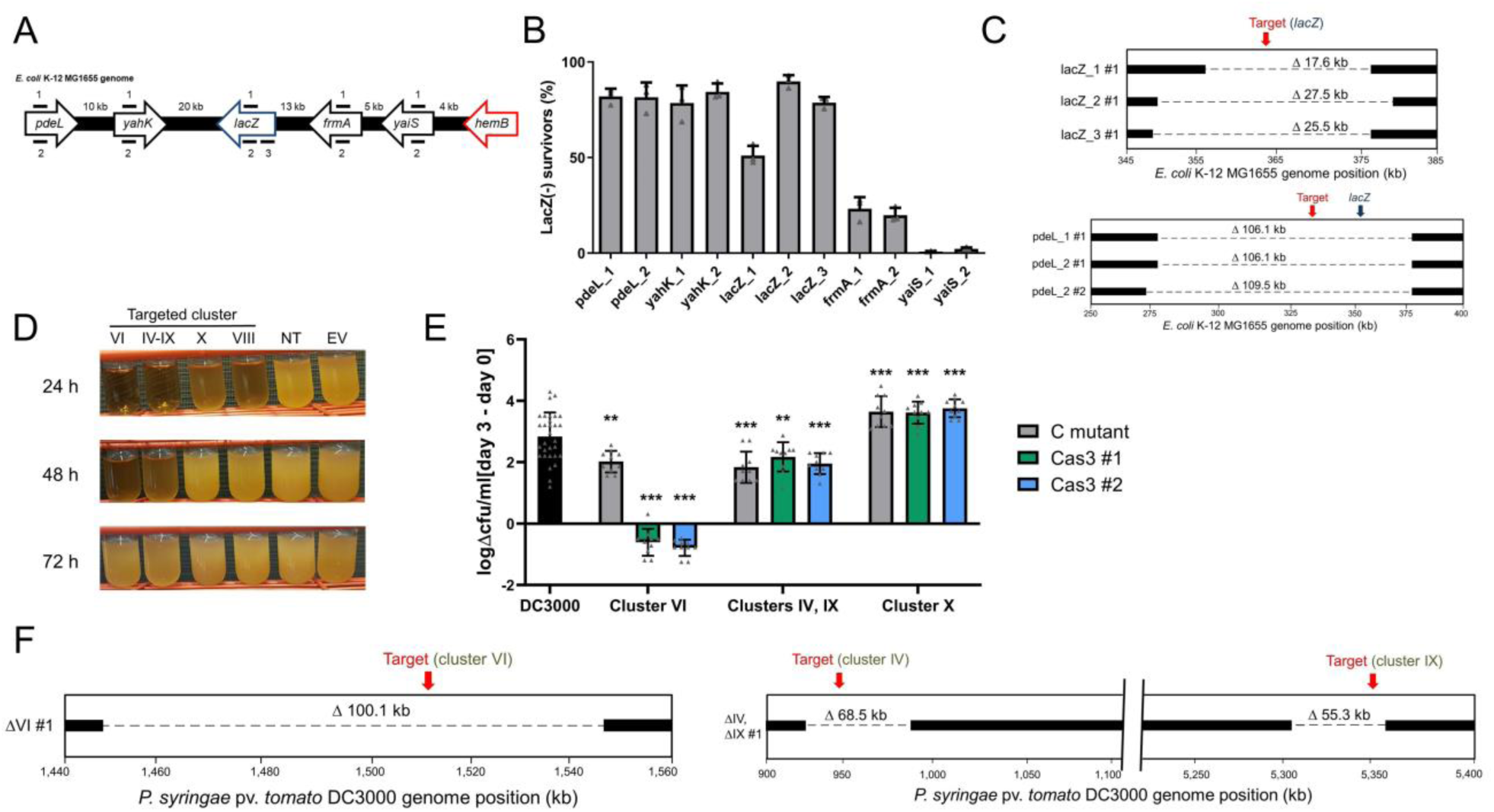
**A)** Schematic of the crRNA targeted sites in the *E. coli* MG1655 genome at the *lacZ* locus. **B)** *lacZ* deletion efficiencies using distinct crRNAs targeting the *E. coil* K-12 MG1655 chromosome. Efficiencies calculated based on LacZ activity. Values are averages of 3 biological replicates, error bars represent standard deviations. **C)** Whole-genome sequencing of an *E. coli* deletion mutant targeted 30 kb upstream of *lacZ* at *pdeL*. **D)** Growth of *P. syringae* DC3000 strains expressing the I-C system and distinct crRNAs. Constructs VI, IV-IX, and VIII target *P. syringae* DC3000 non-essential chromosomal genes, non-targeting crRNA (NT), empty vector (EV). **E)** Bacterial growth of deletion mutants in *Arabidopsis thaliana*. Values are differences in colony forming units (cfu) / ml counted on day 0 of the experiment and day 3, shown on a logarithmic scale. The wild-type DC3000 strain is shown in black, while gray bars represent previously constructed polymutant control (C) strains of the different clusters (labeled at bottom), and green and blue bars show deletion mutants generated using Cas3 (two isolated strains for each targeted cluster, #1, #2). Values shown are means of 10 biological replicates each (30 for DC3000), error bars show SD values, ** p < 0.01, *** p < 0.005, ANOVA analysis (see methods). **F)** Whole-genome sequencing of *P. syringae* deletion mutants. Left panel shows virulence cluster VI targeting, while right panel shows virulence cluster IV and IX targeting with a single crRNA, as the clusters share sequence identity.

Next, we tested Cas3-mediated editing in the plant pathogen *Pseudomonas syringae* pv. *tomato* DC3000, which does not naturally encode a CRISPR-Cas system^30^. *P. syringae* encodes many non-essential virulence effector genes whose activities are difficult to disentangle due to their redundancy^31^. We designed crRNAs targeting four chromosomal virulence effector clusters (IV, VI, VIII, and IX), or one plasmid cluster (pDC3000^32^, cluster X) in *P. syringae* strain DC3000. Two clusters (IV and IX) shared identical sequences that could be targeted simultaneously using a single crRNA. Expression of targeting crRNAs led to a noticeable growth delay compared to non-targeted controls (Figure 4D). PCR analysis of surviving cells showed editing efficiencies of 67-92% (Supplementary Figure 9A). *In planta* and *in vitro* growth assays of three deletion mutants effectively recapitulated the phenotypes of previously described cluster deletion polymutants^32^ (Figure 4E, Supplementary Figure 9B-G). Targeting cluster X cured the 73 kb plasmid and simultaneous cluster IV and IX targeting led to dual deletions in 8 out of 12 survivors, with a sequenced representative having 68.5 kb and 55.3 kb deletions, respectively, at the expected target sites. The effector cluster VI Cas3-derived mutant had a more severe growth defect *in vitro* and *in planta* than the control mutant (Figure 4E, Supplementary Figure 9B,9E). This large deletion (100.1 kb in size) likely impacted general fitness (Figure 4F), demonstrating one drawback of large deletions, but this can be easily overcome by assessing *in vitro* growth of multiple isolates. Using our portable minimal system, we achieved three new applications: the single-step deletion of large virulence regions, multiplexed targeting, and plasmid curing. Overall, we have demonstrated I-C CRISPR-Cas3 editing to be a generally applicable tool capable of generating large genomic deletions in three distinct bacteria.

### Repurposing endogenous CRISPR-Cas3 systems for gene editing

Type I CRISPR-Cas3 systems are the most common CRISPR-Cas systems in nature^1^. Therefore, many bacteria have a built-in genome editing tool to be harnessed. We first tested the environmental isolate from which our Type I-C system was derived. Introduction of the self-targeting *phzM* crRNA led to the isolation of genomic deletions at the targeted site according to PCR analysis in 0-30% of survivors when wild-type repeats flanked the spacer and 30-60% of survivors when modified repeats were used (Supplementary Figure 10A). Whole-genome sequencing of two of these isolates revealed 33.7 (wild-type repeat) and 39 kb (MR) deletions of the target gene and surrounding regions (Figure 5A). Additionally, HDR-based editing with a single construct was again efficacious, with 7/10 survivors acquiring the specific 0.17 kb deletion (Supplementary Figure 10B). Together, these experiments demonstrate the capacity for different forms of genome editing using a single plasmid and an endogenous CRISPR-Cas system.

**Figure 5.**
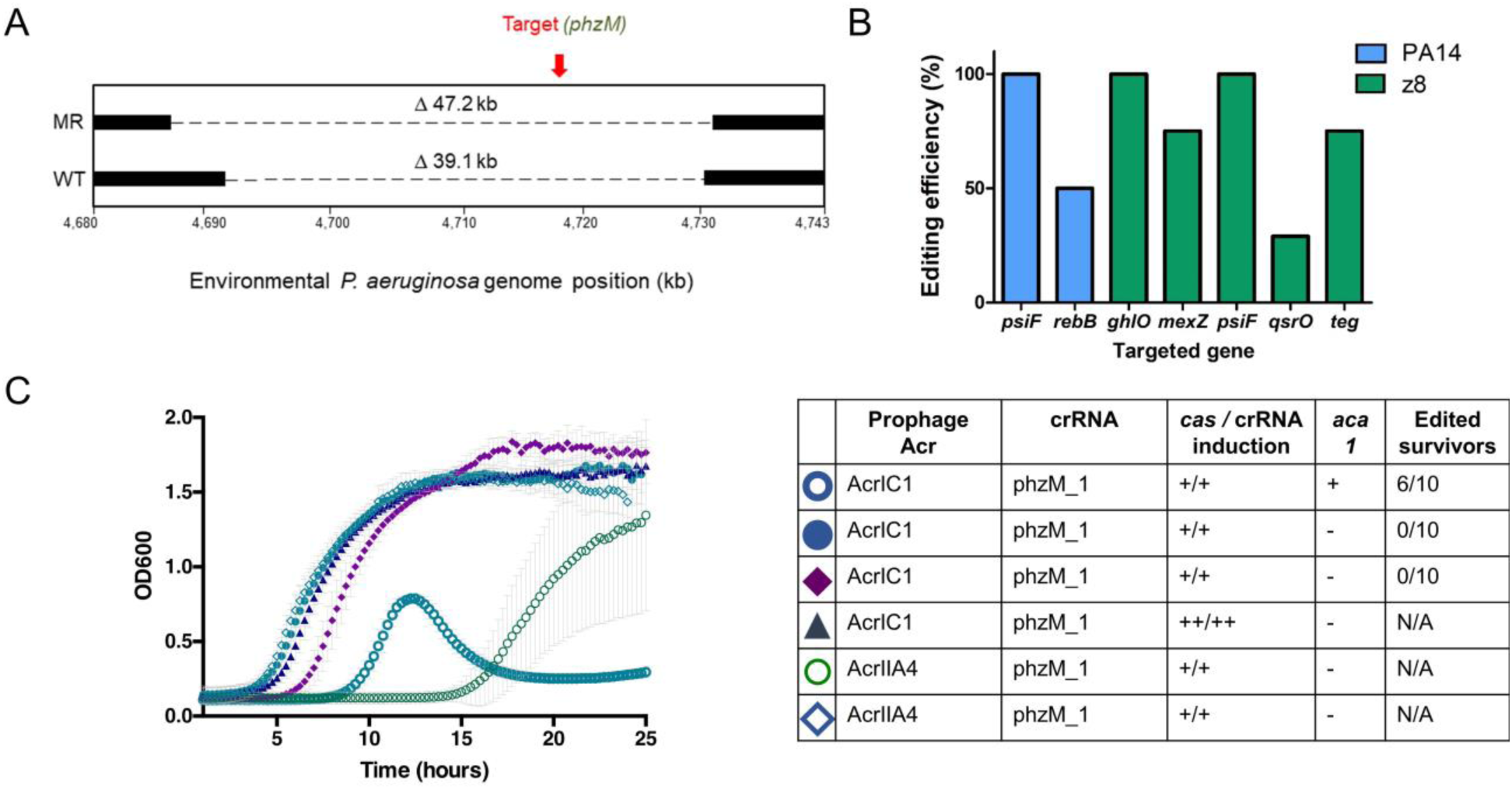
**A)** Schematic of whole genome sequencing of an environmental isolate of PAO1 with an endogenous Type I-C system. Two survivors were isolated post-targeting using either WT direct repeats flanking the spacer, or modified repeats. **B)** Editing efficiencies at targeted genomic sites using homologous templates in a laboratory (PA14) and clinical (z8) strain of *P. aeruginosa*. See Supplementary Table 3 for additional details. **C)** Growth curves of PAO1^IC^ lysogenized by recombinant DMS3m phage expressing *acrIIA4* or *acrIC1* from the native *acr* locus. CRISPR-Cas3 activity is induced with either 0.5mM (+) or 5mM (++) IPTG and 0.1% (+) or 0.3% (++) arabinose. Edited survivors reflect number of isolated survivor colonies missing the targeted gene (*phzM*). Each growth curve is the average of 10 biological replicates and error bars represent SD.

We next evaluated the feasibility of repurposing other Type I systems, using the naturally active Type I-F systems^33^ encoded by laboratory strain *P. aeruginosa* PA14, and the clinical strain *P. aeruginosa* z8. Plasmids with Type I-F specific crRNAs were expressed, targeting various genomic sites for deletion (Supplementary Table 3). HDR templates (600 bp arms on average) were included in the plasmids to generate deletions of defined coordinates ranging from 0.2 to 6.3 kb. Overall, at 5 different genomic target sites in strain z8 and 2 sites in PA14, we observed desired deletions in 29 - 100 % of analyzed survivor colonies (Figure 5B). This demonstrates a high recombination efficiency and the feasibility of repurposing other Type I CRISPR-Cas3 systems in the manner we employed for the minimal I-C system.

Finally, one potential impediment to the implementation of any CRISPR-Cas bacterial genome editing tool is the presence of anti-CRISPR (acr) proteins that inactivate CRISPR-Cas activity^34^. In the presence of a prophage expressing AcrIC1 (a Type I-C anti-CRISPR protein^26^) from a native *acr* promoter, self-targeting was completely inhibited, but not by an isogenic prophage expressing a Cas9 inhibitor AcrIIA4^35^ (Figure 5C). To attempt to overcome this impediment, we expressed *aca1* (anti-CRISPR associated gene 1), a direct negative regulator of *acr* promoters, from the same construct as the crRNA. Using this repression-based “anti-anti-CRISPR” strategy, CRISPR-Cas function was re-activated, allowing the isolation of edited cells despite the presence of *acrIC1* (Figure 5C and Supplementary Figure 10C). In contrast, simply increasing *cas* gene and crRNA expression did not overcome AcrIC1-mediated inhibition (Figure 5C). Therefore, using anti-anti-CRISPRs presents a viable route towards enhanced efficiency of CRISPR-Cas editing and necessitates continued discovery and characterization of anti-CRISPR proteins and their cognate repressors.

## Discussion

By repurposing a minimal CRISPR-Cas3 system as both an endogenous and heterologous genome editing tool, we show that hurdles to generating large deletions can be overcome. We obtained high efficiencies after modifying a repeat sequence to prevent spacer loss. Using only a single crRNA, we isolated deletions as large as 424 kb without requiring the insertion of a selectable marker or HDR templates guiding the repair process. Additionally, the I-C system appears to produce bi-directional deletions, similar to what was previously observed with the I-F CRISPR-Cas3 system^36,37^, but not with type I-E^9,10,22^. CRISPR-Cas3 presents a genome editing tool useful for the targeted removal of large elements (e.g. virulence clusters, plasmids) and also for unbiased screening and genome streamlining. As a long-term goal of microbial gene editing has been genome minimization^38,39,57^, we used our optimized CRISPR-Cas3 system to generate ten iterative deletions, achieving >13 % genome reduction of the targeted strain. This spanned only 30 days while maintaining editing efficiency, a great improvement over previous genome reduction methods^40^. Some basic microbial applications of Cas3 include studying chromosome biology (e.g. replichore asymmetry^41^), virulence factors^42^, and the impact of the mobilome.

An important outcome of this work is the enhanced recombination observed at cut sites when comparing Cas3 and Cas9 directly. The potential for Cas3 to be more recombinogenic through the generation of exposed ssDNA may be advantageous for both programmed knock-outs and for programmed knock-ins. The direct comparison between Cas3 (large deletions) and Cas9 (small deletions) presented here confirms the causality of Cas3 in the deletion outcomes.

Our study has revealed some of the benefits and challenges of working with CRISPR-Cas3. While some of the iteratively edited strains demonstrated slight growth defects, the Cas3 editing workflow shows high potential for genome minimization efforts. Since many distinct deletion events are generated, screening various isolates for fitness benefits or defects is possible, and one can proceed with the strain that has the desired fitness property. Additionally, if the location of essential genes within an organism’s genome is not fully described, this is not an impediment as editing efficiency drops precipitously when targeting near essential genes. Finally, despite our success at transplanting the minimal Type I-C system, it remains to be seen whether the approach will be limited by differences in DNA repair mechanisms. Indeed, in *E. coli* and *P. syringae*, larger regions of homology, such as 34 bp long REP sequences were observed, indicating the role of RecA-mediated homologous recombination^43^ in the repair process. Meanwhile in *P. aeruginosa*, the borders of the deletions showed either small (4-14 bp) micro-homologies or no noticeable sequence homology. The former implies a role for alternative end-joining^29^, while the latter non-homologous end-joining^44^ in the repair process. Downstream studies are required to dissect the roles of each mechanism in the deletion generation process for better predictable deletion outcomes.

CRISPR-Cas3 is an especially promising tool for use in eukaryotic cells as it would facilitate the interrogation of large segments of non-coding DNA, much of which has unknown function^45^. Additionally, it was recently shown that Cas9-generated “gene knockouts” (i.e. small indels causing out-of-frame mutations) frequently encode pseudo-mRNAs that may produce protein products, necessitating methods for full gene removal^46,47^. Encouragingly, a Type I-E CRISPR-Cas system was recently delivered to human cells as a ribonucleoprotein (RNP) complex and resulted in large deletions^22^, demonstrating the feasibility of Cas3-mediated editing in human cells. Overall, the intrinsic properties of Cas3 make it a promising tool to fill a void in current gene editing capabilities. Employing Cas3 to make large genomic deletions will facilitate the manipulation of repetitive and non-coding regions, having a broad impact on genetics research by providing a tool to probe genomes *en masse*, as well as the capability to rapidly streamline genomes for synthetic biology.

## Methods

### Bacterial strains, plasmids, DNA oligonucleotides, and media

A previously described^26^ environmental strain of *Pseudomonas aeruginosa* was used as a template to amplify the *four* cas genes of the Type I-C CRISPR-Cas system genes (*cas3, cas5, cas7*, and *cas8*). The genes were cloned into the pUC18-mini-Tn7T-LAC vector^48^ using the SacI-PstI restriction endonuclease cut sites in the order *cas5, cas7, cas8, cas3* to generate the plasmid pJW31 (Addgene number: 136423). This vector was introduced into *Pseudomonas aeruginosa* PAO1^49^, inserting the *cas* genes into the chromosome, following previously described methods^50^. Following integration, the excess sequences, including the antibiotic resistance marker, were removed via Flp-mediated excision as described previously^50^. The resulting strain, dubbed PAO1^IC^, allowed for inducible expression of the I-C system through induction with isopropyl β-D-1 thiogalactopyranoside (IPTG). An isogenic strain carrying Cas9 derived from *Streptococcus pyogenes* was constructed in the same fashion, resulting in the strain PAO1^IIA^. For experiments to test the system in *Pseudomonas syringae*, we employed the previously characterized strain DC3000^30^. *E. coli* editing experiments were conducted with strain K-12 MG1655^51^.

To achieve genomic self-targeting of the I-C CRISPR-Cas strains, crRNAs designed to target the genome were expressed from the pHERD20T and pHERD30T shuttle vectors^52^. So-called “entry vectors” pHERD20T-ICcr and pHERD30T-ICcr were first generated by cloning at the EcoRI and HindIII sites an annealed linear dsDNA template carrying the I-C CRISPR-Cas system repeat sequences flanking two BsaI Type IIS restriction endonuclease recognition sites. Additionally, a preexisting BsaI site in a non-coding site of the pHERD30T and pHERD20T plasmids was mutated using whole-plasmid amplification so it would not interfere with the cloning of the crRNAs^26^. Oligonucleotides with repeat-specific overhangs encoding the various spacer sequences were annealed and phosphorylated using T4 polynucleotide kinase (PNK) and cloned into the entry vectors using the BsaI sites. For experiments using Cas9, sgRNAs were expressed from the same pHERD30T vector, with the sgRNA construct cloned using the same restriction sites as with the I-C crRNAs.

The all-in-one vector pCas3cRh (Addgene number 133773) is a derivative of the pHERD30T-IC plasmid, with the 4 I-C system genes cloned downstream of the crRNA site. This was achieved by amplifying the genes *cas3, cas5, cas8*, and *cas7* in two fragments with a junction within *cas8* designed to eliminate an intrinsic BsaI site with a synonymous point mutation. The amplified fragments were cloned into pHERD30T-IC using the Gibson assembly protocol^53^. Finally, to guard against potential leaky toxic expression, we replaced the *araC*-Para_BAD_ promoter with the rhamnose-inducible *rhaSR*-Prha_BAD_ system^54^. The sequence for *rhaSR*-Prha_BAD_ was amplified from the pJM230 template^54^, provided by the lab of Joanna B. Goldberg (Emory University), and cloned into the pHERD30T-IC plasmid to replace *araC*-Para_BAD_ using Gibson Assembly (New England Biolabs). Without induction, transformation efficiencies of targeting constructs of assembled pCas3cRh were on average 5-10-fold lower when compared to non-targeting controls (Supplementary Figure 8C), indicating residual leakiness of the I-C system.

The *aca1*-containing vector pICcr-*aca1* is a derivative of the pHERD30T-ICcr plasmid, with *aca1* cloned downstream of the crRNA site under the control of the pBAD promoter. The *aca1* gene was cloned from *P. aeruginosa* phage DMS3m.

All oligonucleotides used in this study were obtained from Integrated DNA Technologies. For a complete list of all DNA oligonucleotides and a short description, see Supplementary Table 4.

*P. aeruginosa* and *E. coli* strains were grown in standard Lysogeny Broth (LB): 10 g tryptone, 5 g yeast extract, and 10 g NaCl per 1 L dH_2_O. Solid plates were supplemented with 1.5 % agar. *P. syringae* was grown in King’s medium B (KB): 20 g Bacto Proteose Peptone No. 3, 1.5 g K_2_HPO_4_, 1.5 g MgSO_4_•7H_2_O, 10 ml glycerol per 1 L dH_2_O, supplemented with 100 μg/ml rifampicin. The following antibiotic concentrations were used for selection: 50 μg/ml gentamicin for *P. aeruginosa* and *P. syringae*, 15 μg/ml for *E. coli*; 50 μg/ml carbenicillin for all organisms. Inducer concentrations were 0.5 mM IPTG, 0.1 % arabinose, and 0.1 % rhamnose. For transformation protocols, all bacteria were recovered in Super optimal broth with catabolite repression (SOC): 20 g tryptone, 5 g yeast extract, 10mM NaCl, 2.5 mM KCl, 10 mM MgCl_2_, 10 mM, MgSO_4_, and 20 mM glucose in 1 L dH_2_O.

### Bacterial transformations

Transformations of *P. aeruginosa, E. coli*, and *P. syringae* strains were conducted using standard electroporation protocols. 10 ml of overnight cultures were centrifuged and washed twice in an equal volume of 300 mM sucrose (20 % glycerol for *E. coli*) and suspended in 1 ml 300 mM sucrose (20 % glycerol for *E. coli*). 100 μl aliquots of the resulting competent cells were electroporated using a Gene Pulser Xcell Electroporation System (Bio-Rad) with 50 – 200 ng plasmid with the following settings: 200 Ω, 25 μF, 1.8 kV, using 0.2 mm gap width electroporation cuvettes (Bio-Rad). Electroporated cells were incubated in antibiotic-free SOC media for 1 hour at 37 °C (28 °C for *P. syringae*), then plated onto LB agar (KB agar for *P. syringae*) with the selecting antibiotic, and grown overnight at 37 °C (28 °C for *P. syringae*). Cloning procedures were performed in commercial *E. coli* DH5α cells (New England Biolabs) or *E. coli* XL1-Blue (QB3 Macrolab Berkeley), according to the manufacturer’s protocols.

### Construction of recombinant DMS3m acr phages

The isogenic DMS3m *acrIIA4* and *acrIC1* phages were constructed using previously described methods^55^. A recombination cassette, pJZ01, was constructed with homology to the DMS3m *acr* locus. Using Gibson Assembly (New England Biolabs), either *acrIC1* or *acrIIA4* were cloned upstream of *aca1*, and the resulting vectors were used to transform PAO1^IC^. The transformed strains were infected with WT DMS3m, and recombinant phages were screened for. Phages were stored in SM buffer at 4 °C.

### Isolation of PAO1^IC^ lysogens

PAO1^IC^ was grown overnight at 37 °C in LB media. 150 μl of overnight culture was added to 4 ml of 0.7 % LB top agar and spread on 1.5 % LB agar plates supplemented with 10 mM MgSO_4_. 5 μl of phage, expressing either *acrIC1* or *acrIIA4* were spotted on the solidified top agar and plates were incubated at 30 °C overnight. Following incubation, bacterial growth within the plaque was isolated and spread on 1.5 % LB agar plate. After an overnight incubation at 37 °C, single colonies were assayed for the prophage. Confirmed lysogens were used for genomic targeting experiments.

### Genomic targeting

#### Pseudomonas aeruginosa

Genomic self-targeting of *P. aeruginosa* PAO1^IC^ was achieved by electroporating cells with pHERD30T (or pHERD20T) expressing the self-targeting spacer of choice. Cells were plated onto LB agar plates containing the selective antibiotic, without inducers, and grown overnight. Single colonies were then grown in liquid LB media containing the selective antibiotic, as well as IPTG to induce the genomic expression of the I-C system genes, and arabinose to induce the expression of the crRNA from the plasmid. The *aca1*-containing crRNA plasmids do not need additional inducers, as the pBAD promoter controls *aca1*. Cultures were grown at 37 °C in a shaking incubator overnight to saturation, then plated onto LB agar plates containing the selecting antibiotic, as well as the inducers, and incubated overnight again at 37 °C. The resulting colonies were then analyzed individually using colony PCR for any differences at the targeted genomic site compared to a wild-type cell. gDNA was isolated by resuspending 1 colony in 20 μl of H_2_O, followed by incubation at 95 °C for 15 min. 1-2 μl of boiled sample was used for PCR. The primers used to assay the targeted sites were designed to amplify genomic regions 1.5 - 3 kb in size. In the event of a PCR product equal to or smaller than the wild-type fragment (as was often observed when analyzing Cas9-targeted cells), Sanger sequencing (Quintara Biosciences) was used to determine any modifications of the targeted sequences. In some cases, additional analysis of the crRNA-expressing plasmids of the surviving colonies was also performed, by isolating and reintroducing the plasmids into the original I-C CRISPR-Cas strain, where functional self-targeting could be determined based on a significant increase in the lag time of induced cultures, characteristic of self-targeting events.

#### Escherichia coli

Genomic self-targeting of *E. coli* was conducted in a similar fashion as *P. aeruginosa*, except using the pCas3cRh all-in-one vector. Electrocompetent *E. coli* cells were transformed with pCas3cRh expressing a crRNA targeting the genome. Individual transformants were selected and grown in liquid LB media containing the selecting antibiotic (gentamicin) overnight without any inducers added. The overnight cultures were then plated in the presence of inducer and X-gal to screen for functional *lacZ* (LB agar + 15 μg/ml gentamicin + 0.1 % rhamnose + 1 mM IPTG + 20 μg/ml X-gal) and blue/white colonies were counted the next day.

#### Pseudomonas syringae

Electrocompetent *P. syringae* cells were also transformed with pCas3cRh plasmids targeting selected genomic sequences. Initial transformants were plated onto KB agar + 100 μg/ml rifampicin + 50 μg/ml gentamicin plates, and incubated at 28 °C overnight. Single colony transformants were then selected and inoculated in KB liquid media supplemented with rifampicin, gentamicin, and 0.1 % rhamnose inducer, and grown to saturation in a shaking incubator at 28 °C. Cultures were finally plated onto KB agar plates with rifampicin, gentamicin, and rhamnose and incubated at 28 °C. Individual colonies were finally assayed with colony PCR to determine the presence of deletions at the targeted genomic sites.

#### Iterative genome minimization

Iterative targeting to generate multiple deletions in the *P. aeruginosa* PAO1^IC^ strain was carried out by alternating the pHERD30T and pHERD20T plasmids each expressing different crRNAs targeting the genome. Each crRNA designed to target the genome was cloned into both the pHERD30T plasmid, which confers gentamicin resistance, as well as the pHERD20T plasmid, which confers carbenicillin resistance. After first transforming and targeting with a pHERD30T plasmid expressing a specific crRNA, deletion candidate isolates were transformed with a pHERD20T expressing a crRNA targeting a different genomic region. As the two plasmids are identical with the exception of the resistance marker, this eliminated the necessity for curing of the original plasmid to be able to target a different region. For the next targeting event, the pHERD30T plasmid could again be used, this time expressing another crRNA targeting a different genomic region. In this manner, pHERD30T and pHERD20T could be alternated to achieve multiple deletions in a rapid process. At each new transformation step, the cells were checked for any residual resistance to the given antibiotic from a previous cycle. Additionally, functionality of the CRISPR-Cas system of the edited cells could be determined through the introduction of a plasmid expressing crRNA targeting the D3 bacteriophage^34^, then performing a phage spotting assay to see if phage targeting was occurring or not.

### Measurement of growth rates

#### Pseudomonas aeruginosa

Growth dynamics of various strains were measured using a Synergy 2 automated 96-well plate reader (Biotek Instruments) and the accompanying Gen5 software (Biotek Instruments). Individual colonies were picked and grown overnight in 300 μl volumes of LB in 96-well deep-well plates at 37 °C. The grown cultures were then diluted 100-fold into 100 μl of fresh LB in a 96-well clear microtitre plate (Costar) and sealed with Microplate sealing adhesive (Thermo Scientific). Small holes were punched in the sealing adhesive for each well for increased aeration. Doubling times were calculated as described previously^56^.

#### Pseudomonas syringae

To test bacterial growth *in planta*, we used the *Arabidopsis thaliana* ecotype *Columbia* (Col-0), which has previously been shown to be susceptible to infection by *P. syringae* DC3000. Plants were grown for 5-6 weeks in 9h light/15h darkness and 65 % humidity. For each inoculum, we measured bacterial growth in 10 individual Col-0 plants. Four leaves from each plant were infiltrated at OD_600_ = 0.0002, and cored with a #3 borer. The four cores from each plant were then ground, resuspended in 10 mM MgCl_2_ and plated in a dilution series on selective media for colony counts at both the time of infection and 3 days post-infection.

To test bacterial growth *in vitro*, we used both KB and plant apoplast mimicking minimal media (MM)^57^. Overnight cultures were prepared from single colonies of each strain, washed, and diluted to OD_600_ = 0.1 in 96-well plates using either KB or MM. Plates were incubated with shaking at 28 °C. OD_600_ was measured over the course of 24-25 hours using an Infinate 200 Pro automated plate reader (Tecan). Statistical analysis determined significantly different groups based on ANOVA analysis on the day 0 group of values and the day 3 group of values. Significant ANOVA results (p<0.01) were further analyzed with a Tukey’s HSD post hoc test to generate adjusted p-values for each pairwise comparison. A significance threshold of 0.01 was used to determine which treatment groups were significantly different.

#### Bacteriophage plaque (spot) assays

Bacteriophage plaque assays were performed using 1.5 % LB agar plates supplemented with 10 mM MgSO_4_ and the appropriate antibiotic (gentamicin or carbenicillin, depending on the plasmid used to express the crRNA), and 0.7 % LB top agar supplemented with 0.5 mM IPTG and 0.1 % arabinose inducers added covering the whole plate. 150 μl of the appropriate overnight cultures was suspended in 4 ml molten top agar poured onto an LB agar plate leading to the growth of a bacterial lawn. After 10-15 minutes at room temperature, 3 μl of ten-fold serial dilutions of bacteriophage was spotted onto the solidified top agar. Plates were incubated overnight at 30 °C and imaged the following day using a Gel Doc EZ Gel Documentation System (BioRad) and Image Lab (BioRad) software. The following bacteriophage were used in this study: bacteriophage JBD30^34^, bacteriophage D3^58^, and bacteriophage DMS3m^59^.

#### Whole-genome sequencing

Genomic DNA for whole-genome sequencing (WGS) analysis was isolated directly from bacterial colonies using the Nextera DNA Flex Microbial Colony Extraction kit (Illumina) according to the manufacturer’s protocol. Genomic DNA concentration of the samples was determined using a DS-11 Series Spectrophotometer/Fluorometer (DeNovix) and all fell into the range of 200-500 ng/μl. Library preparation for WGS analysis was done using the Nextera DNA Flex Library Prep kit (Illumina) according to the manufacturer’s protocol starting from the tagment genomic DNA step. Tagmented DNA was amplified using Nextera DNA CD Indexes (Illumina). Samples were placed overnight at 4 °C following the tagmented DNA amplification step, then continued the next day with the library clean up steps. Quality control of the pooled libraries was performed using a 2100 Bioanalyzer Instrument (Agilent Technologies) with a High Sensitivity DNA Kit (Agilent Technologies). The majority of samples were sequenced using a MiSeq Reagent Kit v2 (Illumina) for a 150 bp paired-end sequencing run using the MiSeq sequencer (Illumina). *P. syringae* and Cas9-generated *P. aeruginosa* deletion strains were sequenced using a NextSeq 500 Reagent Kit v2 (Illumina) for a 150 bp paired-end sequencing run using the NextSeq 500 sequencer (Illumina).

Genome sequence assembly was performed using Geneious Prime software version 2019.1.3. Paired read data sets were trimmed using the BBDuk (Decontamination Using Kmers) plugin using a minimum Q value of 20. The genome for the ancestral PAO1^IC^ strain was de novo assembled using the default automated sensitivity settings offered by the software. The consensus sequence of PAO1^IC^ assembled in this manner was then used as the reference sequence for mapping all of the PAO1^IC^ strains with multiple deletions. As a control, the sequences were also mapped to the reference *P. aeruginosa* PAO1 sequence (NC_002516) to verify deletion border coordinates. Coverage of these sequenced strains ranged from 66 to 143-fold, with an average of 98.3-fold. The sequenced *P. aeruginosa* environmental strains were also mapped to the PAO1 (NC_002516) reference, while the sequenced *E. coli* strains were mapped to the *E. coli* K-12 MG1655 reference sequence (NC_000913). Finally, sequenced *P. syringae* strains were mapped to the *P. syringae* DC3000 (NC_004578) reference sequence, along with the pDC3000A endogenous 73.5 kb plasmid sequence (NC_004633). All of the remaining sequenced strains had > 100-fold coverage. All deletion junction sequences were manually verified by the presence of multiple reads spanning the deletions, containing sequences from both end boundaries.

WGS data was visualized using the BLAST Ring Image Generator (BRIG) tool^60^ employing BLAST+ version 2.9.0. In several cases, short sequences were aligned within previously determined large deletions at redundant sequences such as transposase genes. Such misrepresentations created by BRIG were manually removed to reflect the actual sequencing data.

## Data availability

Raw whole-genome sequencing data associated with Figures 1D, 3B, 4A, 4C, 4F, and 5A) has been uploaded to GenBank (Submission number SUB6598604, accession number pending) and is also available, along with bacterial strains, upon request from the corresponding author.

## Acknowledgements

We thank Joanna B. Goldberg (Emory University) for providing the plasmid pJM230, and Adair Borges (UCSF) for providing pAB04 to clone Type I-F crRNAs. We thank the Bondy-Denomy lab for productive conversations pertaining to this project.

## Funding

B.C. is supported by the Eötvös National Scholarship of Hungary and a Marie Skłodowska-Curie Actions Individual Global Fellowship (number 844093) of the Horizon 2020 Research Program of the European Commission. L.L. is supported by the HHMI Gilliam Fellowship for Advanced Study and the UCSF Discovery Fellowship. Research on plant immunity in the Lewis laboratory is supported by the USDA ARS 2030-21000-046-00D and 2030-21000-050-00D (J.D.L.), and the NSF Directorate for Biological Sciences IOS-1557661 (J.D.L.). I.J.C. is supported by a Grace Kase fellowship from UC Berkeley and the NSF Graduate Research Fellowship Program. A.V.R. is supported by funding from the Pew Charitable Trusts. E.D.C. is funded by the Chan Zuckerberg Biohub. CRISPR-Cas3 projects in the Bondy-Denomy Lab are supported by the UCSF Program for Breakthrough Biomedical Research funded in part by the Sandler Foundation and an NIH Director’s Early Independence Award DP5-OD021344.

## Competing Interests

J.B.-D. is a scientific advisory board member of SNIPR Biome and Excision Biotherapeutics and a scientific advisory board member and co-founder of Acrigen Biosciences.

---

**Supplementary Figure 1.**
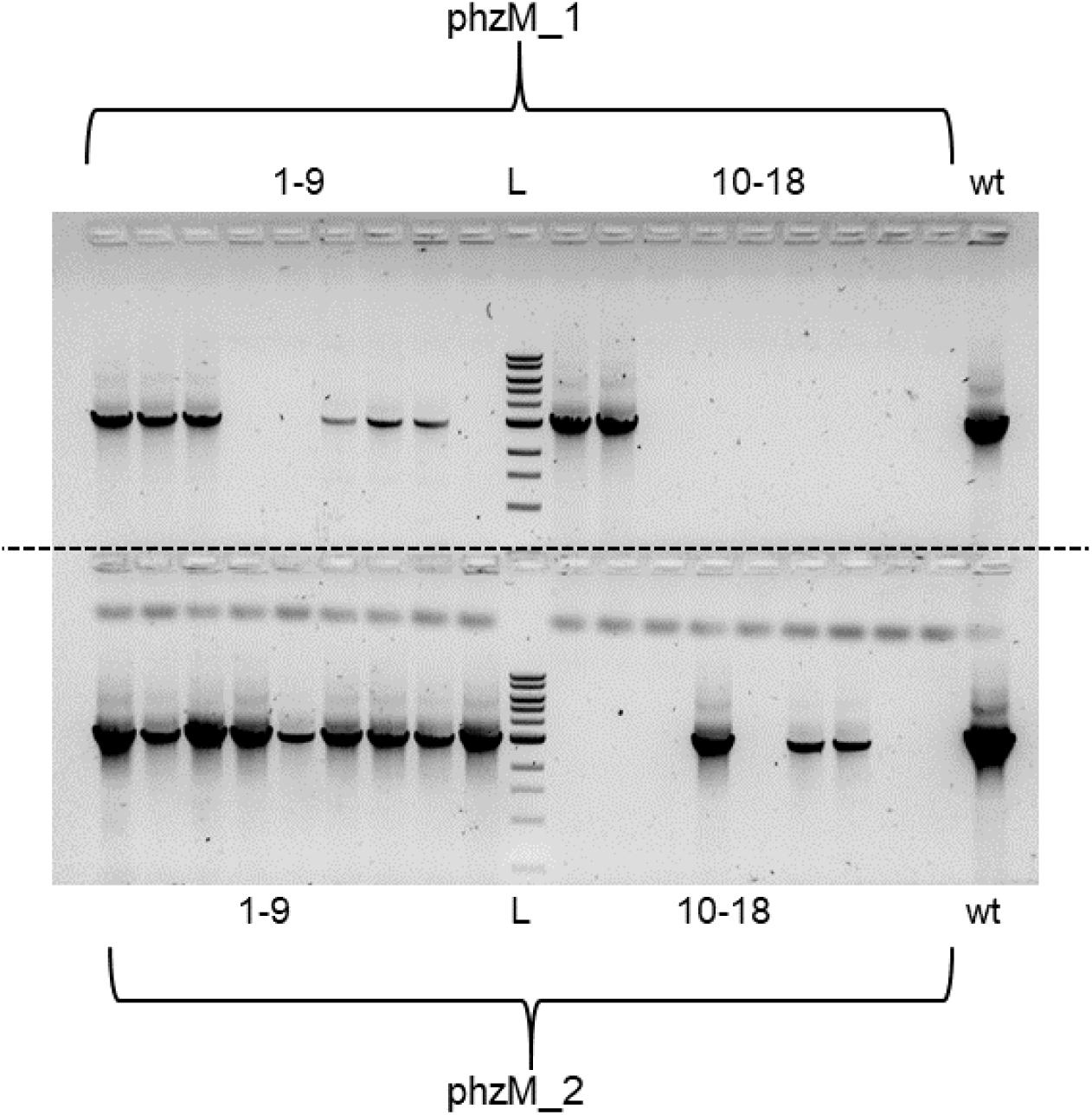
PCR amplification of a 3 kb genomic fragment flanking the *phzM* gene targeted using two different crRNAs, phzM_1 and phzM_2. Colony PCRs were performed on 18 biological replicates of self-targeted strains for each crRNA. The PAO1^IC^ parental strain is used as a positive control (wt). L indicates a 1 kb DNA ladder.

**Supplementary Figure 2.**
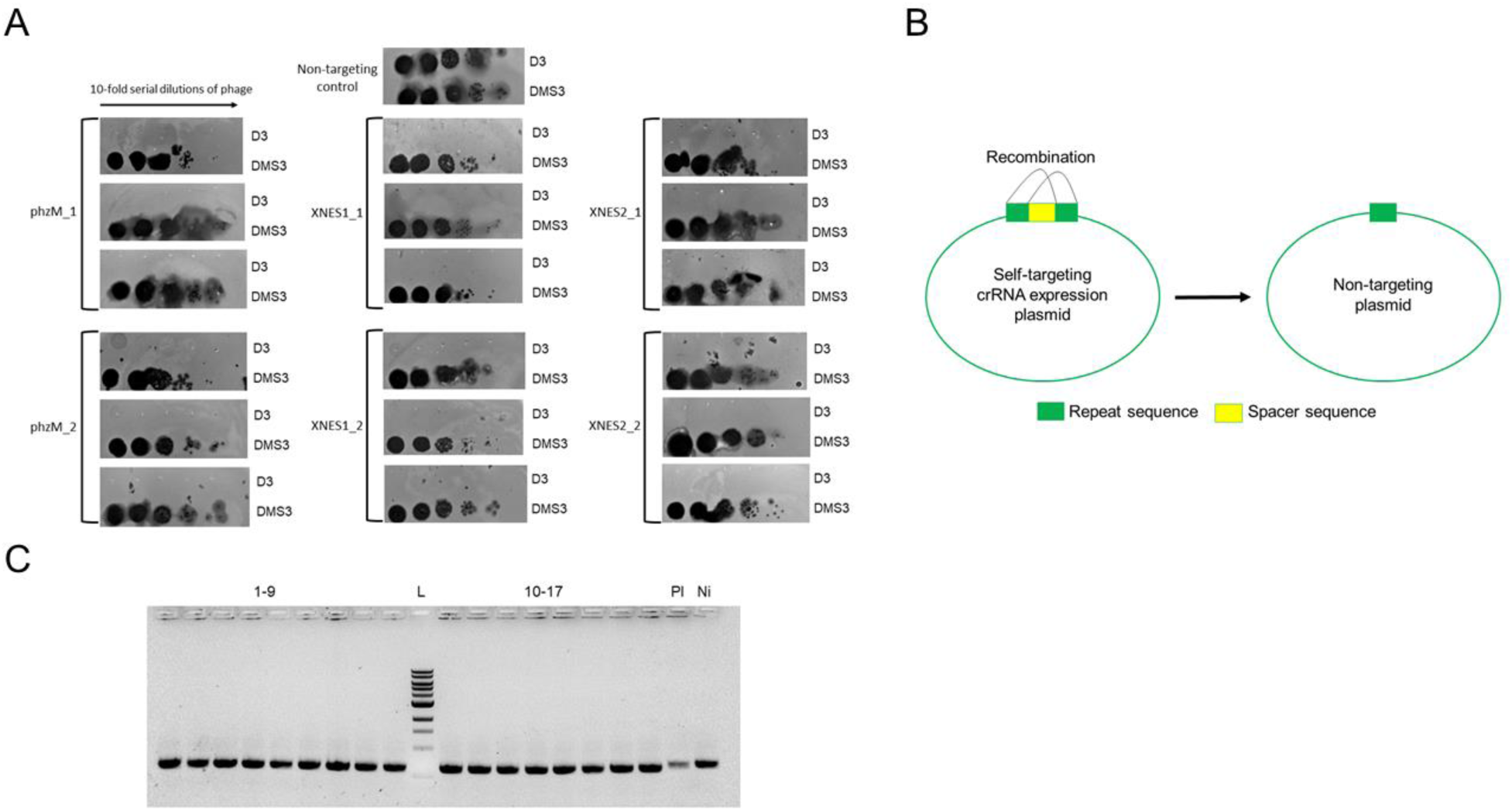
**A)** Phage targeting assays with survivors that had no discernable deletion of the crRNA-targeted genomic site. Strains were transformed with a D3 phage-targeting crRNA to assay for IC CRISPR-Cas3 activity. Three unique survivors were isolated from six self-targeting assays for a total of 18 survivors. Control is a non-targeting crRNA. **B)** Schematic of spacer excision events where the two direct repeats recombine, resulting the loss of the targeting spacer. **C)** PCR amplification of the crRNA sequence from plasmids isolated from 17 non-deletion self-targeted survivors. Pl indicates the original plasmid as the PCR template, Ni indicates a sample where the crRNA was not induced, L indicates a 1 kb DNA ladder.

**Supplementary Figure 3.**
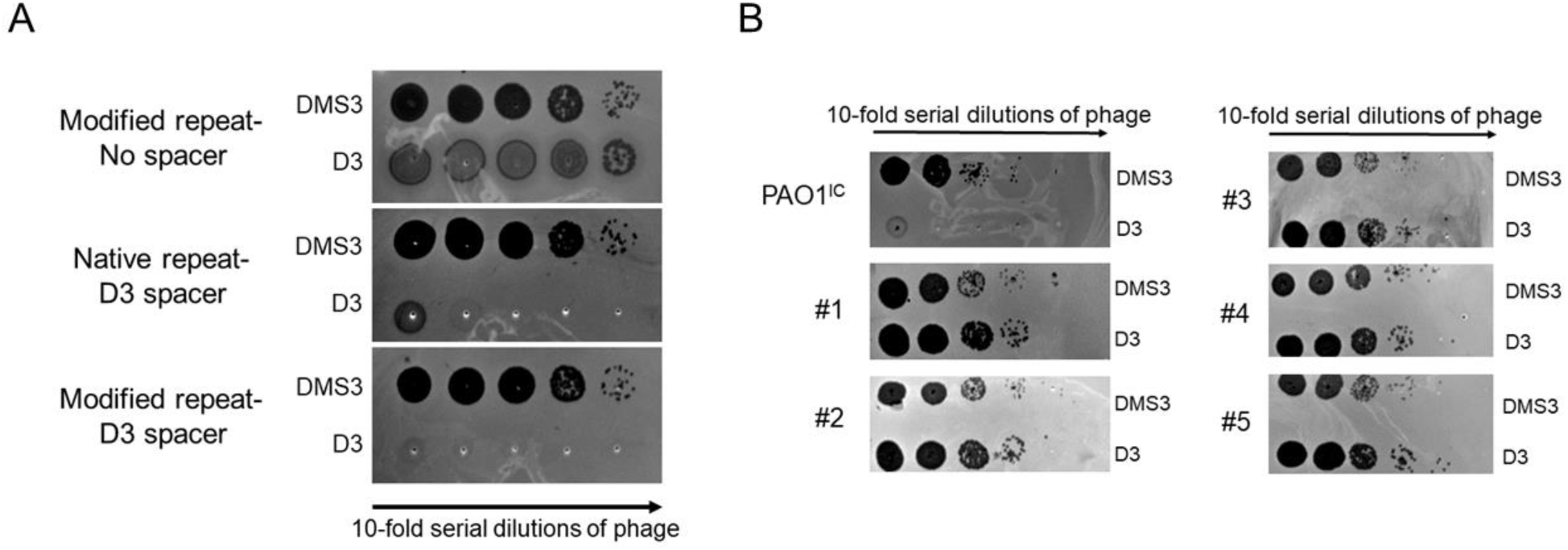
**A)** Phage-targeting assay showing the activity of the modified repeat crRNA constructs. Ten-fold serial dilutions of DMS3 phage and D3 phage were spotted on lawns of PAO1^IC^ expressing either empty vector (top), a crRNA targeting D3 with WT direct repeats (middle), or a crRNA targeting D3 with modified repeats (bottom). **B)** Phage targeting assay of five non-deletion self-targeting survivors expressing a D3 phage targeting crRNA. Unsuccessful targeting of phage indicates a non-functional CRISPR-Cas system in these strains. The parental PAO1^IC^ strain with a functional CRISPR-Cas system was used as a control.

**Supplementary Figure 4.**
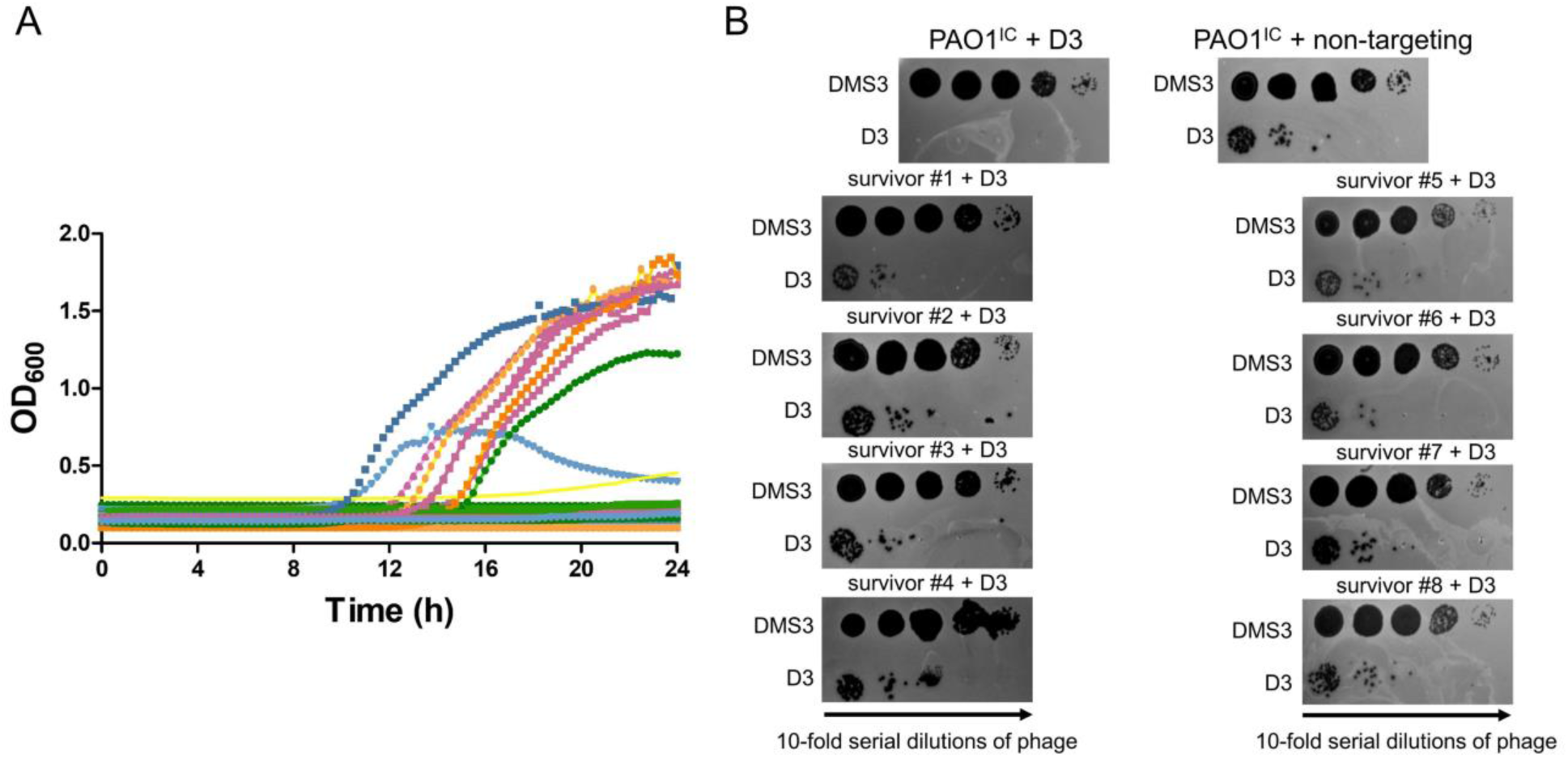
**A)** Growth curves of 36 PAO1^IC^ biological replicates targeting the essential gene, *rplQ*, using the MR crRNA plasmid. **B)** Phage targeting assays with eight isolated *rplQ*-targeted survivors to assay for I-C CRISPR-Cas activity. Serial dilutions of DMS3 phage and D3 phage were spotted on lawns of PAO1^IC^ expressing a crRNA targeting phage D3. The parent PAO1^IC^ strain expressing a D3 targeting crRNA (top left) was used as a positive control, while PAO1^IC^ expressing a non-targeting crRNA was used as a negative control.

**Supplementary Figure 5.**
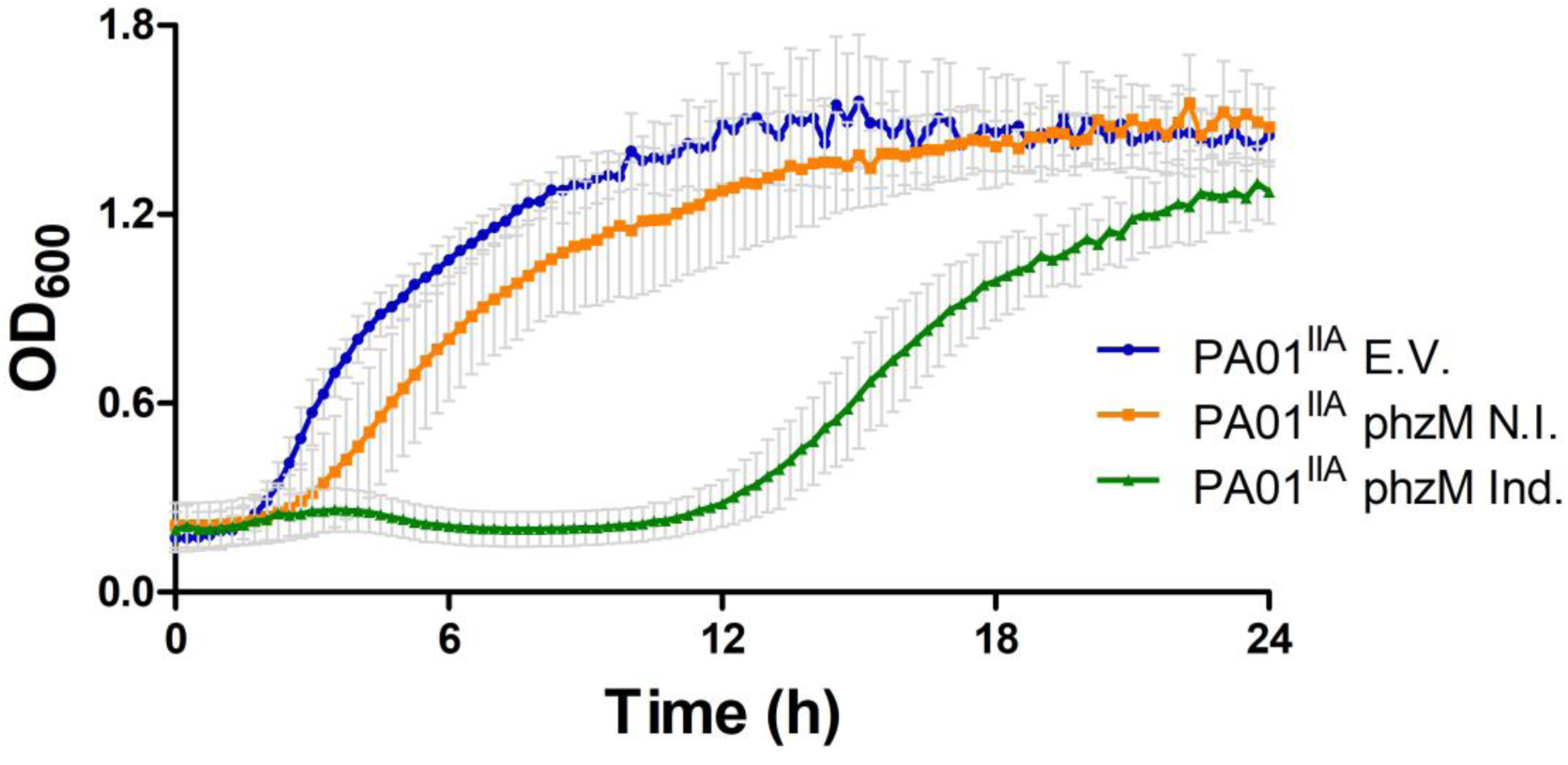
Growth of self-targeting strains of PAO1^IIA^ expressing a self-targeting crRNA targeting the genome at *phzM* (Ind.). An empty vector (E.V.) and a non-induced *phzM* targeting strain (N.I.) were used as controls. Mean OD values measured at 600 nm are shown for 8 biological replicates each, error bars indicate SD values.

**Supplementary Figure 6.**
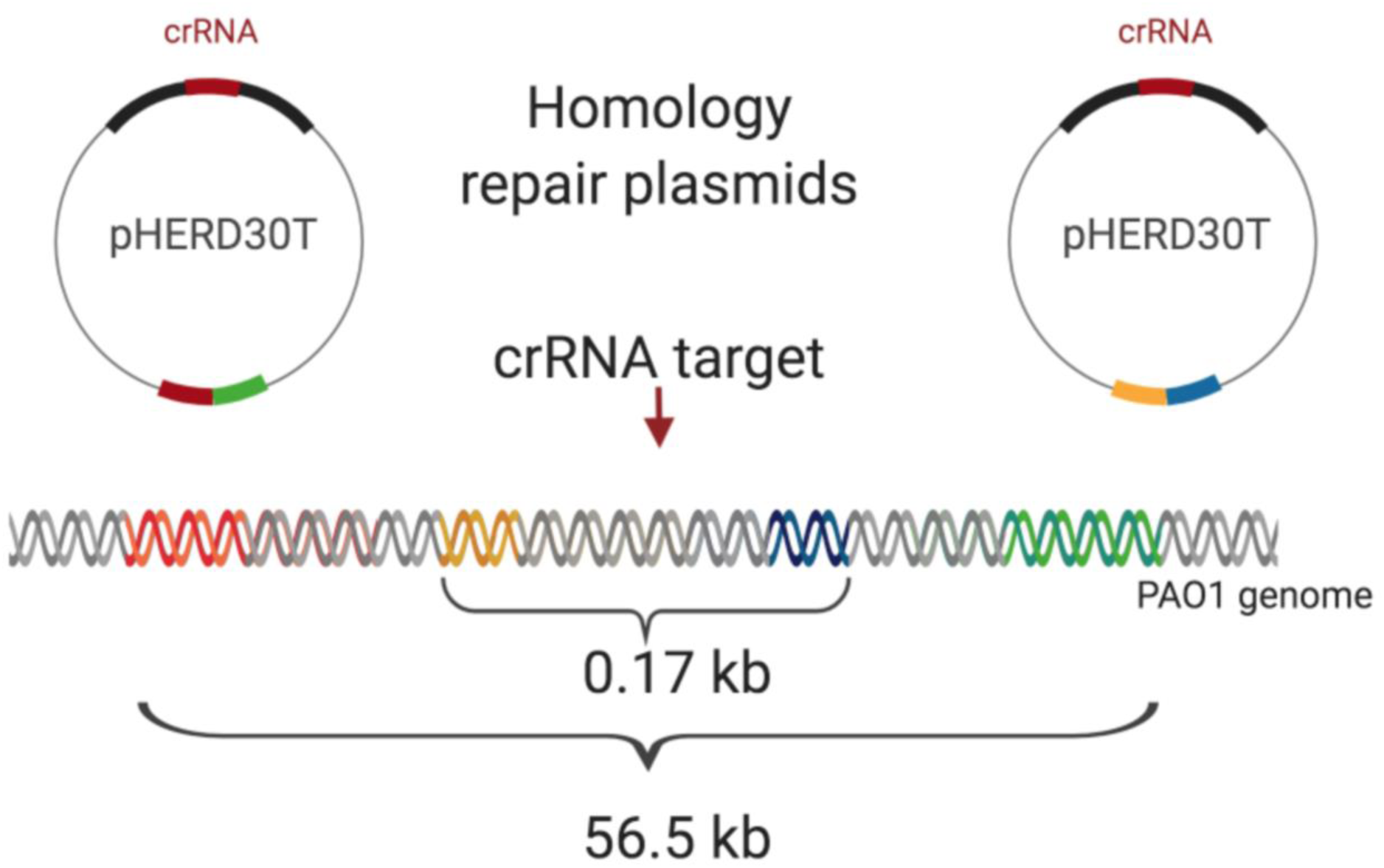
Schematic overview of the generation of deletions with predetermined coordinates of various sizes. Sequences with ∼400 bp homology to genomic sites (purple and yellow boxes for the short deletion, red and orange boxes for the long deletion) were cloned into the vector crRNA vector.

**Supplementary Figure 7.**
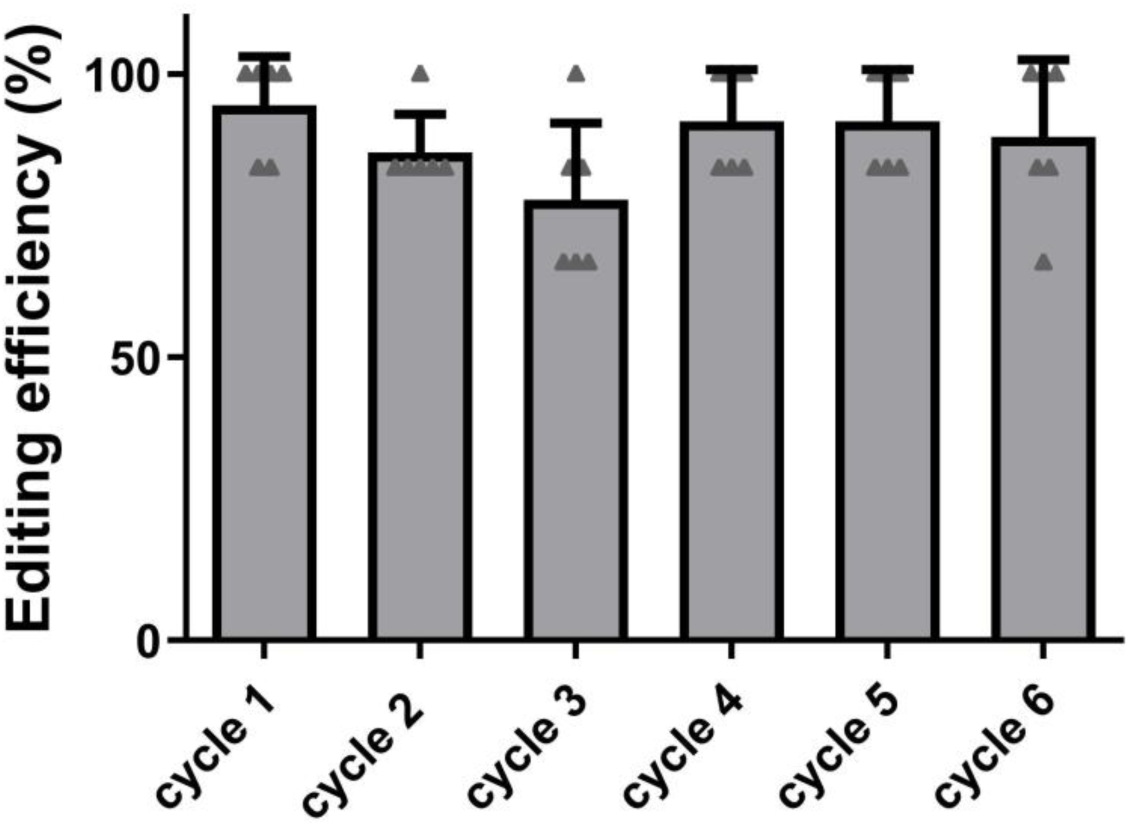
Deletion efficiencies observed over six cycles of iterative self-targeting. Six genomic targets were targeted in six different orders. Six survivors were analyzed using site-specific PCR after each cycle, for a total of 36 analyzed colonies (6*6) after each cycle.

**Supplementary Figure 8.**
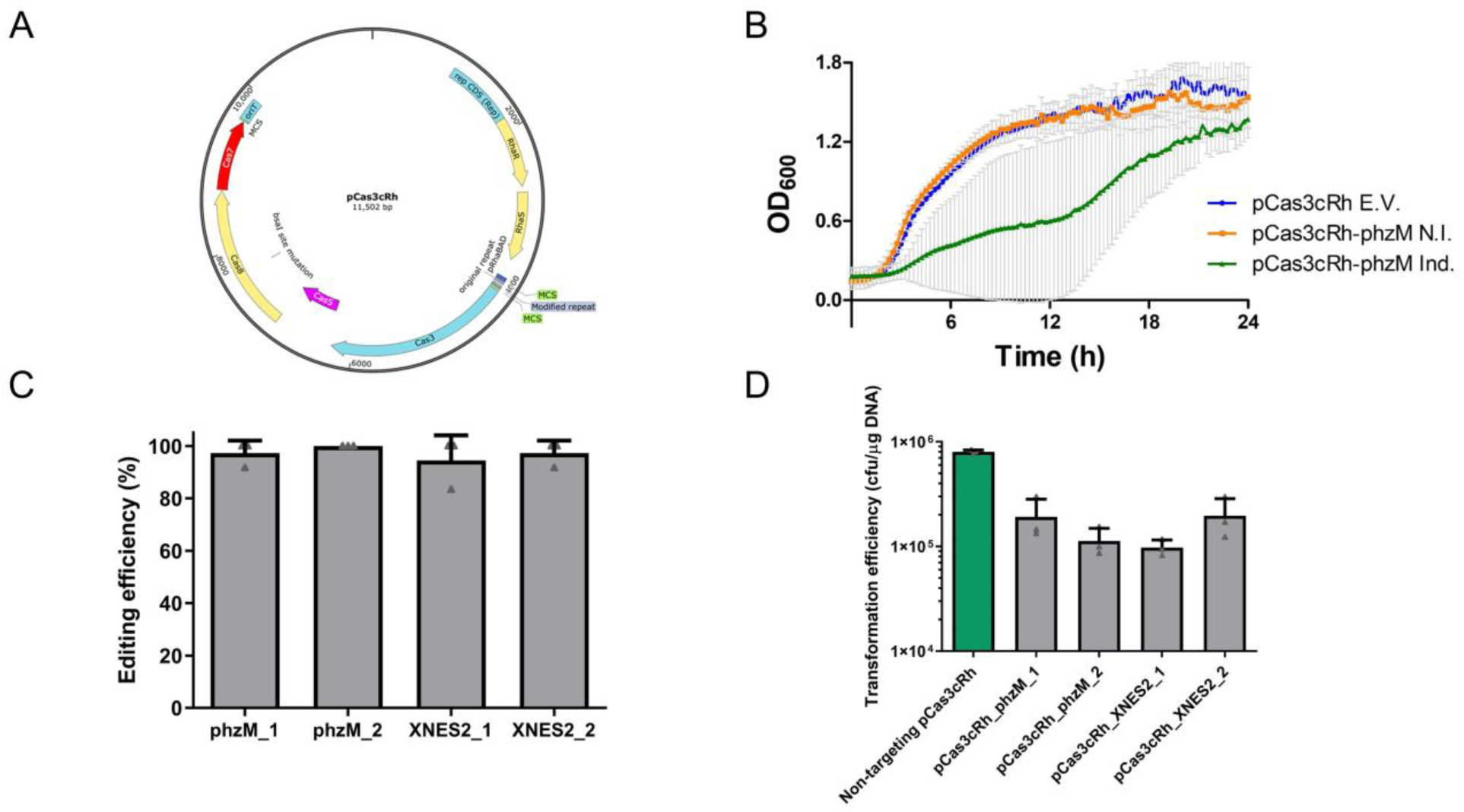
**A)** Map of the I-C CRISPR-Cas all-in-one plasmid pCas3cRh carrying I-C crRNA and genes *cas3, cas5, cas8*, and *cas7* under the control of the rhamnose-inducible *rhaSR*-Prha_BAD_ system. **B)** Growth curve of PAO1 transformed with the pCas3cRh vector expressing a self-targeting crRNA targeting *phzM* (Ind.). An empty vector (E.V.) and a non-induced *phzM* targeting strain (N.I.) were used as controls. Mean OD values measured at 600 nm are shown for six biological replicates each. **C)** Deletion efficiencies for WT PAO1 using the all-in-one vector pCas3cRh carrying all necessary components of the I-C CRISPR-Cas system. Values are averages of three replicates where 12 individual colonies were analyzed using site-specific PCR. Error bars show standard deviations. **D)** Transformation efficiencies with self-targeting pCas3cRh vectors expressing crRNAs for *phzM* or XNES 2 compared to a non-targeting control (green bar) in PAO1. Values are means of 3 replicates each, error bars represent SD values.

**Supplementary Figure 9.**
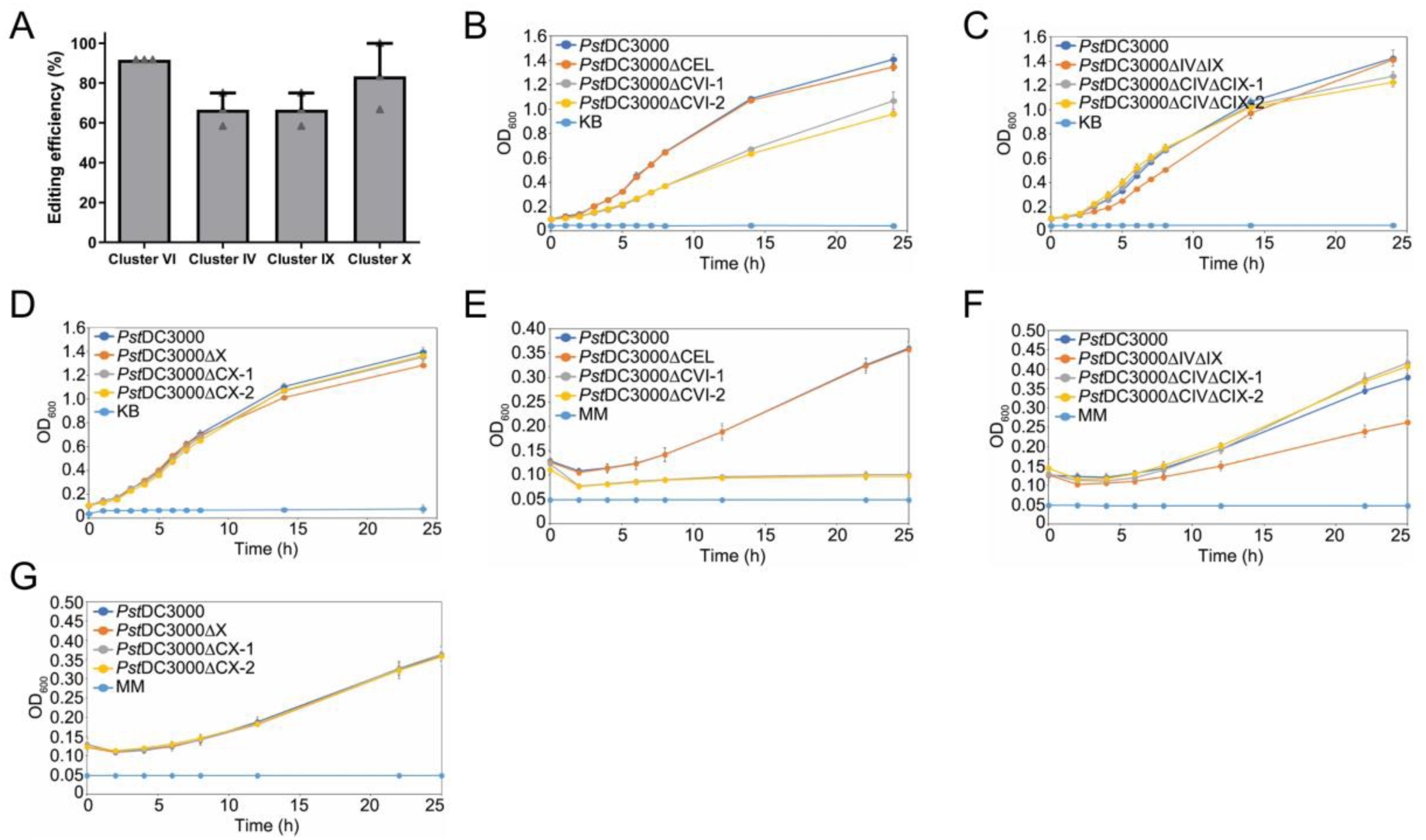
**A)** Percentage of survivors with targeted deletions in clusters of non-essential virulence effector genes in *P. syringae* pv. *tomato* DC3000. Values are averages of three biological replicates where 12 individual colonies were analyzed using site-specific PCR for each, error bars show standard deviations. **B)** *In vitro* growth of cluster VI deletion strains in King’s medium B (KB). ΔCEL is the previously published polymutant, while ΔCVI-1 and ΔCVI-2 are Cas3-generated mutants. Error bars represent standard deviation, n = 4. **C)** *In vitro* growth of cluster IV, cluster IX deletion strains in KB. ΔCEL is the previously published polymutant, while ΔCIVΔCIX-1 and ΔCIVΔCIX-2 are Cas3-generated mutants. Error bars represent standard deviation, n = 4. **D)** *In vitro* growth of cluster X deletion strains in KB. ΔCEL is the previously published polymutant, while ΔCX-1 and ΔCX-2 are Cas3-generated mutants. Error bars represent standard deviation, n = 4. **E)** *In vitro* growth of cluster VI deletion strains in apoplast mimicking minimal media (MM). ΔCEL is the previously published polymutant, while ΔCVI-1 and ΔCVI-2 are Cas3-generated mutants. Error bars represent standard deviation, n = 4. **F)** *In vitro* growth of cluster IV, cluster IX deletion strains in MM. ΔCEL is the previously published polymutant, while ΔCIVΔCIX-1 and ΔCIVΔCIX-2 are Cas3-generated mutants. Error bars represent standard deviation, n = 4. **G)** *In vitro* growth of cluster X deletion strains in MM. ΔCEL is the previously published polymutant, while ΔCX-1 and ΔCX-2 are Cas3-generated mutants. Error bars represent standard deviation, n = 4.

**Supplementary Figure 10.**
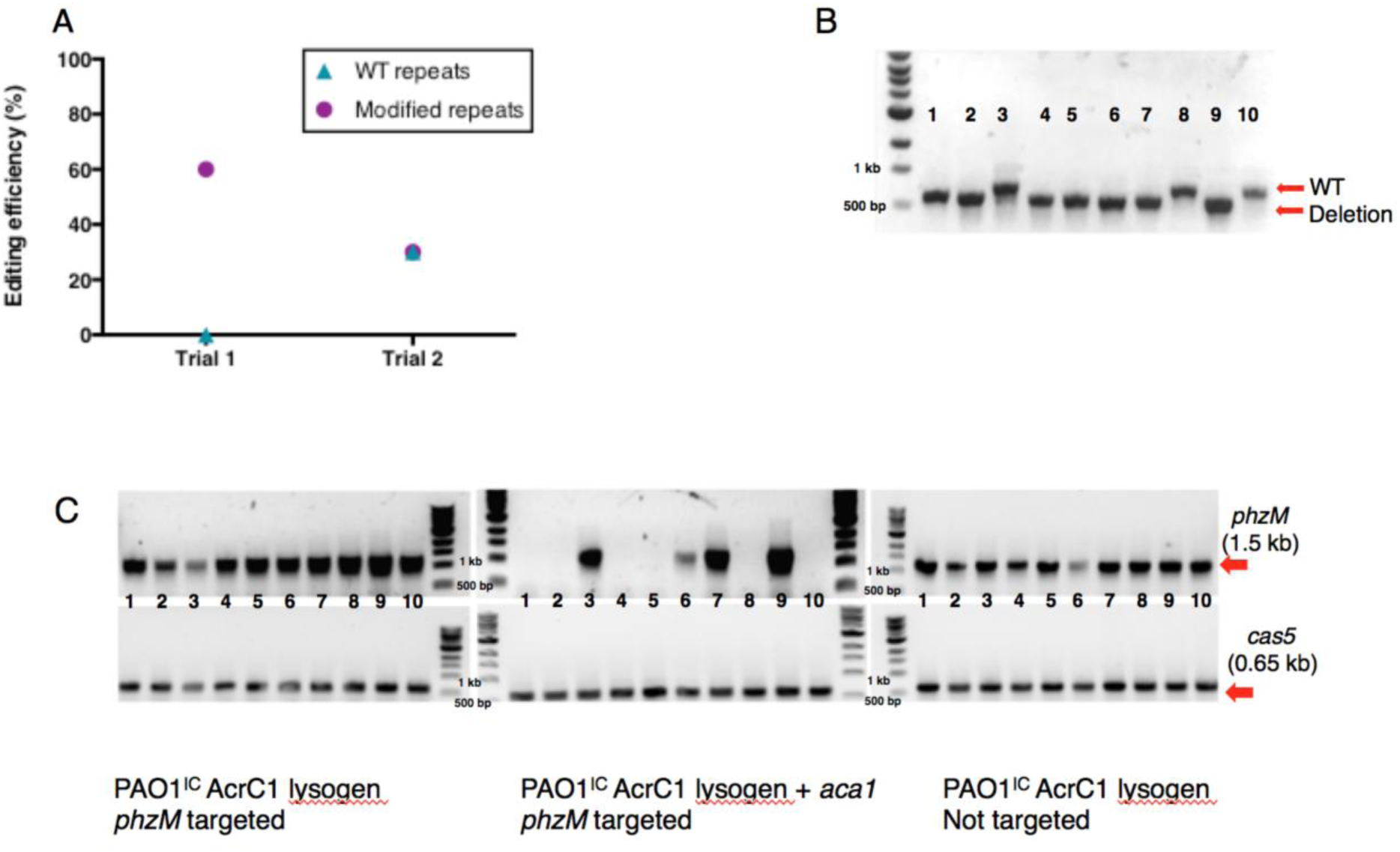
**A)** Editing efficiencies for the *Pseudomonas aeruginosa* environmental isolate naturally expressing the Type I-C *cas* genes, transformed with a plasmid targeting *phzM* with WT repeats or modified repeats. Each data point represents the fraction of isolates with the deletion out of ten isolates assayed. **B)** Genotyping results for the *Pseudomonas aeruginosa* environmental isolate using the 0.17 kb HDR template. Larger band corresponds to the WT sequence, smaller band corresponds to a genome reduced by 0.17 kb. **C)** Genotyping results of PAO1^IC^ AcrC1 lysogens after self-targeting induction in the presence or absence of *aca1* and a non-targeted control. Ten biological replicates per strain were assayed. gDNA was extracted from each replicate and PCR analysis for the *phzM* gene (targeted gene, top row of gels) or *cas5* gene (non-targeted gene, bottom row) was conducted. Only cells that co-expressed *aca1* with the crRNA showed loss of the *phzM* band, indicating genome editing. All replicates had a *cas5* band, indicating successful gDNA extraction and target specificity for the *phzM* locus.

**Supplementary Table 1.** E<u>x</u>tended, <u>n</u>on-<u>es</u>sential regions (XNES) of *P. aeruginosa* PAO1 genome with contiguous, individually non-essential genes in a complex laboratory medium exceeding 100 kb. Data based on a transposon sequencing dataset from Turner *et al*.^27^.

| Region | Coordinates | Size |
| --- | --- | --- |
| XNES 1 | 27535 – 142359 | 114 kb |
| XNES 2 | 143267 – 371151 | 228 kb |
| XNES 3 | 491900 – 606160 | 114 kb |
| XNES 4 | 841825 – 986817 | 145 kb |
| XNES 5 | 1147815 – 1249907 | 102 kb |
| XNES 6 | 1260442 – 1491913 | 232 kb |
| XNES 7 | 1974210 – 2150828 | 176 kb |
| XNES 8 | 2216121 – 2375804 | 160 kb |
| XNES 9 | 2376541 – 2923367 | 546 kb |
| XNES 10 | 2972700 – 3079197 | 106 kb |
| XNES 11 | 3155072 – 3309411 | 154 kb |
| XNES 12 | 3587303 – 3802567 | 216 kb |
| XNES 13 | 3897357 – 4062426 | 165 kb |
| XNES 14 | 4294208 – 4457362 | 163 kb |
| XNES 15 | 4576324 – 4753990 | 178 kb |
| XNES 16 | 6025305 – 6180942 | 156 kb |

**Supplementary Table 2.** Genomic coordinates and extent of homologous sequences at genomic deletion junctions of whole-genome sequenced self-targeting strains of *P. aeruginosa, P. syringae*, and *E. coli*. See separate Excel File.

**Supplementary Table 3.** Summary of HR-mediated genome editing experiments using the Type I-F CRISPR-Cas3 system. Genes were targeted for deletion in the strains PA14 and z8. Experiments targeted 4 single genes and 2 gene blocks, *teg* and *rebB*, that comprise X and Y genes, respectively. Transformants were classified as 1) ‘HR edits’ that have the HR designed deletion; 2) ‘non-templated edits’ that have a non-designed deletion encompassing the targeted gene, 3) ‘no edits’ where the targeted gene is intact. (*) two colony morphologies with different editing frequencies were obtained in this experiment.

| Strain | Edited gene | HR edits (%) | Nontemplated edits (%) | No edits (%) | n | Designed deletion (bp) | HR template length (left + right, bp) |
| --- | --- | --- | --- | --- | --- | --- | --- |
| PA14 | *psiF* | 100 | 0 | 0 | 12 | 0.5 | 600 + 600 |
| PA14 | *rebB* | 50 | 50 | 0 | 16 | 4.1 | 600 + 600 |
| z8 | *ghlO* | 100 | 0 | 0 | 5 | 0.2 | 600 + 600 |
| z8 | *mexZ* | 75 | 25 | 0 | 12 | 0.6 | 722 + 600 |
| z8 | *psiF* | 100 | 0 | 0 | 12 | 0.5 | 600 + 600 |
| z8 | *qsrO* | 29(80)* | 71 | 0 | 14 | 0.4 | 751 + 596 |
| z8 | *teg* | 75 | 25 | 0 | 4 | 6.3 | 800 + 809 |

**Supplementary Table 4.** List of oligonucleotides (including crRNA sequences) used in the study. See separate Excel File.

## References

1. Makarova, K. S. et al. An updated evolutionary classification of CRISPR-Cas systems. Nat. Rev. Microbiol. 13, 722–736 (2015).

2. Barrangou, R. et al. CRISPR provides acquired resistance against viruses in prokaryotes. Science 315, 1709–1712 (2007).

3. Garneau, J. E. et al. The CRISPR/Cas bacterial immune system cleaves bacteriophage and plasmid DNA. Nature 468, 67–71 (2010).

4. Barrangou, R. & Doudna, J. A. Applications of CRISPR technologies in research and beyond. Nat. Biotechnol. 933–941 (2016) doi:10.1038/nbt.3659.

5. Wiedenheft, B. et al. Structures of the RNA-guided surveillance complex from a bacterial immune system. Nature 477, 486–489 (2011).

6. Westra, E. R. et al. CRISPR immunity relies on the consecutive binding and degradation of negatively supercoiled invader DNA by Cascade and Cas3. Mol. Cell 46, 595–605 (2012).

7. Brouns, S. J. J. et al. Small CRISPR RNAs Guide Antiviral Defense in Prokaryotes. Science 321, 960–964 (2008).

8. Sinkunas, T. et al. Cas3 is a single-stranded DNA nuclease and ATP-dependent helicase in the CRISPR/Cas immune system. EMBO J. 30, 1335–1342 (2011).

9. Sinkunas, T. et al. In vitro reconstitution of Cascade-mediated CRISPR immunity in Streptococcus thermophilus. EMBO J. 32, 385–394 (2013).

10. Mulepati, S. & Bailey, S. In vitro reconstitution of an Escherichia coli RNA-guided immune system reveals unidirectional, ATP-dependent degradation of DNA target. J. Biol. Chem. 288, 22184–22192 (2013).

11. Hochstrasser, M. L. et al. CasA mediates Cas3-catalyzed target degradation during CRISPR RNA-guided interference. Proc. Natl. Acad. Sci. U. S. A. 111, 6618–6623 (2014).

12. Redding, S. et al. Surveillance and Processing of Foreign DNA by the Escherichia coli CRISPR-Cas System. Cell 163, 854–865 (2015).

13. Xiao, Y., Luo, M., Dolan, A. E., Liao, M. & Ke, A. Structure basis for RNA-guided DNA degradation by Cascade and Cas3. Science 361, eaat0839 (2018).

14. Esvelt, K. M. & Wang, H. H. Genome-scale engineering for systems and synthetic biology. Mol. Syst. Biol. 9, (2013).

15. Montalbano, A., Canver, M. C. & Sanjana, N. E. High-Throughput Approaches to Pinpoint Function within the Noncoding Genome. Mol. Cell 68, 44–59 (2017).

16. Vercoe, R. B. et al. Cytotoxic Chromosomal Targeting by CRISPR/Cas Systems Can Reshape Bacterial Genomes and Expel or Remodel Pathogenicity Islands. PLOS Genet 9, e1003454 (2013).

17. Li, Y. et al. Harnessing Type I and Type III CRISPR-Cas systems for genome editing. Nucleic Acids Res. 44, e34–e34 (2016).

18. Pyne, M. E., Bruder, M. R., Moo-Young, M., Chung, D. A. & Chou, C. P. Harnessing heterologous and endogenous CRISPR-Cas machineries for efficient markerless genome editing in *Clostridium*. Sci. Rep. 6, 25666 (2016).

19. Zhang, J., Zong, W., Hong, W., Zhang, Z.-T. & Wang, Y. Exploiting endogenous CRISPR-Cas system for multiplex genome editing in Clostridium tyrobutyricum and engineer the strain for high-level butanol production. Metab. Eng. doi:10.1016/j.ymben.2018.03.007.

20. Maikova, A., Kreis, V., Boutserin, A., Severinov, K. & Soutourina, O. Using endogenous CRISPR-Cas system for genome editing in the human pathogen Clostridium difficile. Appl. Environ. Microbiol. AEM.01416-19 (2019) doi:10.1128/AEM.01416-19.

21. Hidalgo-Cantabrana, C., Goh, Y. J., Pan, M., Sanozky-Dawes, R. & Barrangou, R. Genome editing using the endogenous type I CRISPR-Cas system in Lactobacillus crispatus. Proc. Natl. Acad. Sci. U. S. A. 116, 15774–15783 (2019).

22. Dolan, A. E. et al. Introducing a Spectrum of Long-Range Genomic Deletions in Human Embryonic Stem Cells Using Type I CRISPR-Cas. Mol. Cell 74, 936-950.e5 (2019).

23. Pickar-Oliver, A. et al. Targeted transcriptional modulation with type I CRISPR–Cas systems in human cells. Nat. Biotechnol. 1–9 (2019) doi:10.1038/s41587-019-0235-7.

24. Nam, K. H. et al. Cas5d Protein Processes Pre-crRNA and Assembles into a Cascade-like Interference Complex in Subtype I-C/Dvulg CRISPR-Cas System. Structure 20, 1574–1584 (2012).

25. Hochstrasser, M. L., Taylor, D. W., Kornfeld, J. E., Nogales, E. & Doudna, J. A. DNA Targeting by a Minimal CRISPR RNA-Guided Cascade. Mol. Cell 63, 840–851 (2016).

26. Marino, N. D. et al. Discovery of widespread type I and type V CRISPR-Cas inhibitors. Science 362, 240–242 (2018).

27. Turner, K. H., Wessel, A. K., Palmer, G. C., Murray, J. L. & Whiteley, M. Essential genome of Pseudomonas aeruginosa in cystic fibrosis sputum. Proc. Natl. Acad. Sci. 112, 4110–4115 (2015).

28. Meek, D. W. & Hayward, R. S. Nucleotide sequence of the rpoA-rplQ DNA of Escherichia coli: a second regulatory binding site for protein S4? Nucleic Acids Res. 12, 5813–5821 (1984).

29. Chayot, R., Montagne, B., Mazel, D. & Ricchetti, M. An end-joining repair mechanism in Escherichia coli. Proc. Natl. Acad. Sci. 107, 2141–2146 (2010).

30. Buell, C. R. et al. The complete genome sequence of the Arabidopsis and tomato pathogen Pseudomonas syringae pv. tomato DC3000. Proc. Natl. Acad. Sci. U. S. A. 100, 10181–10186 (2003).

31. Lindeberg, M., Cunnac, S. & Collmer, A. Pseudomonas syringae type III effector repertoires: last words in endless arguments. Trends Microbiol. 20, 199–208 (2012).

32. Kvitko, B. H. et al. Deletions in the repertoire of Pseudomonas syringae pv. tomato DC3000 type III secretion effector genes reveal functional overlap among effectors. PLoS Pathog. 5, e1000388 (2009).

33. Cady, K. C., Bondy-Denomy, J., Heussler, G. E., Davidson, A. R. & O’Toole, G. A. The CRISPR/Cas adaptive immune system of Pseudomonas aeruginosa mediates resistance to naturally occurring and engineered phages. J. Bacteriol. 194, 5728–5738 (2012).

34. Bondy-Denomy, J., Pawluk, A., Maxwell, K. L. & Davidson, A. R. Bacteriophage genes that inactivate the CRISPR/Cas bacterial immune system. Nature 493, 429–432 (2013).

35. Rauch, B. J. et al. Inhibition of CRISPR-Cas9 with Bacteriophage Proteins. Cell 168, 150-158.e10 (2017).

36. Richter, C. et al. Priming in the Type I-F CRISPR-Cas system triggers strandindependent spacer acquisition, bi-directionally from the primed protospacer. Nucleic Acids Res. 42, 8516–8526 (2014).

37. Rollins, M. F. et al. Cas1 and the Csy complex are opposing regulators of Cas2/3 nuclease activity. Proc. Natl. Acad. Sci. U. S. A. 114, E5113–E5121 (2017).

38. Pósfai, G. et al. Emergent properties of reduced-genome Escherichia coli. Science 312, 1044–1046 (2006).

39. Fehér, T., Papp, B., Pál, C. & Pósfai, G. Systematic Genome Reductions: Theoretical and Experimental Approaches. Chem. Rev. 107, 3498–3513 (2007).

40. Csörgő, B., Nyerges, Á., Pósfai, G. & Fehér, T. System-level genome editing in microbes. Curr. Opin. Microbiol. 33, 113–122 (2016).

41. Képès, F. et al. The layout of a bacterial genome. FEBS Lett. 586, 2043–2048 (2012).

42. Ghosh, S. & O’Connor, T. J. Beyond Paralogs: The Multiple Layers of Redundancy in Bacterial Pathogenesis. Front. Cell. Infect. Microbiol. 7, (2017).

43. Kowalczykowski, S. C. & Eggleston, A. K. Homologous Pairing and Dna Strand-Exchange Proteins. Annu. Rev. Biochem. 63, 991–1043 (1994).

44. Bowater, R. & Doherty, A. J. Making Ends Meet: Repairing Breaks in Bacterial DNA by Non-Homologous End-Joining. PLOS Genet 2, e8 (2006).

45. Hnisz, D. et al. Super-enhancers in the control of cell identity and disease. Cell 155, 934–947 (2013).

46. Tuladhar, R. et al. CRISPR-Cas9-based mutagenesis frequently provokes on-target mRNA misregulation. Nat. Commun. 10, 1–10 (2019).

47. Smits, A. H. et al. Biological plasticity rescues target activity in CRISPR knock outs. Nat. Methods 1–7 (2019) doi:10.1038/s41592-019-0614-5.

48. Choi, K.-H. et al. A Tn7-based broad-range bacterial cloning and expression system. Nat. Methods 2, 443–448 (2005).

49. Stover, C. K. et al. Complete genome sequence of Pseudomonas aeruginosa PAO1, an opportunistic pathogen. Nature 406, 959 (2000).

50. Choi, K.-H. & Schweizer, H. P. mini-Tn7 insertion in bacteria with single attTn7 sites: example Pseudomonas aeruginosa. Nat. Protoc. 1, 153–161 (2006).

51. Blattner, F. R. et al. The complete genome sequence of Escherichia coli K-12. Science 277, 1453–1462 (1997).

52. Qiu, D., Damron, F. H., Mima, T., Schweizer, H. P. & Yu, H. D. PBAD-Based Shuttle Vectors for Functional Analysis of Toxic and Highly Regulated Genes in Pseudomonas and Burkholderia spp. and Other Bacteria. Appl. Environ. Microbiol. 74, 7422–7426 (2008).

53. Gibson, D. G. et al. Enzymatic assembly of DNA molecules up to several hundred kilobases. Nat. Methods 6, 343–345 (2009).

54. Meisner, J. & Goldberg, J. B. The Escherichia coli rhaSR-PrhaBAD Inducible Promoter System Allows Tightly Controlled Gene Expression over a Wide Range in Pseudomonas aeruginosa. Appl. Environ. Microbiol. 82, 6715–6727 (2016).

55. Borges, A. L. et al. Bacteriophage Cooperation Suppresses CRISPR-Cas3 and Cas9 Immunity. Cell 174, 917-925.e10 (2018).

56. Nyerges, Á. et al. Directed evolution of multiple genomic loci allows the prediction of antibiotic resistance. Proc. Natl. Acad. Sci. 115, E5726–E5735 (2018).

57. Huynh, T. V., Dahlbeck, D. & Staskawicz, B. J. Bacterial blight of soybean: regulation of a pathogen gene determining host cultivar specificity. Science 245, 1374–1377 (1989).

58. Kropinski, A. M. Sequence of the Genome of the Temperate, Serotype-Converting, Pseudomonas aeruginosa Bacteriophage D3. J. Bacteriol. 182, 6066–6074 (2000).

59. Budzik, J. M., Rosche, W. A., Rietsch, A. & O’Toole, G. A. Isolation and Characterization of a Generalized Transducing Phage for Pseudomonas aeruginosa Strains PAO1 and PA14. J. Bacteriol. 186, 3270–3273 (2004).

60. Alikhan, N.-F., Petty, N. K., Ben Zakour, N. L. & Beatson, S. A. BLAST Ring Image Generator (BRIG): simple prokaryote genome comparisons. BMC Genomics 12, 402 (2011).

